# Extracellular Matrix Alterations in Chronic Ischemic Cardiomyopathy Revealed by Quantitative Proteomics

**DOI:** 10.1101/2025.05.23.655833

**Authors:** Kevin M. Buck, Holden T. Rogers, Zachery R. Gregorich, Morgan W. Mann, Timothy J. Aballo, Zhan Gao, Emily A. Chapman, Andrew J. Perciaccante, Scott J. Price, Ienglam Lei, Paul C. Tang, Ying Ge

**Affiliations:** Department of Chemistry, University of Wisconsin-Madison, Madison, Wisconsin, 53706, USA; Department of Cell and Regenerative Biology, University of Wisconsin-Madison, Madison, Wisconsin, 53705, USA; Department of Animal and Dairy Sciences, University of Wisconsin-Madison, Madison, Wisconsin, 53706, USA; Department of Medicine, University of Wisconsin-Madison, Madison, WI, USA; Molecular and Cellular Pharmacology Training Program, University of Wisconsin-Madison, Madison, WI 53705, USA; Department of Physiology and Biomedical Engineering, Mayo Clinic, Rochester, Minnesota 55905, USA; Department of Cardiac Surgery, Mayo Clinic, Rochester, Minnesota, 55905, USA; Human Proteomics Program, School of Medicine and Public Health, University of Wisconsin-Madison, Madison, Wisconsin, 53705, USA

## Abstract

Ischemic cardiomyopathy (ICM) is a leading cause of heart failure characterized by extensive remodeling of the cardiac extracellular matrix (ECM). While initially adaptive, ECM deposition following ischemic injury eventually turns maladaptive, promoting adverse cardiac remodeling. The strong link between the extent of fibrosis and adverse clinical outcomes has led to growing interest in ECM targeted therapies to prevent or reverse maladaptive cardiac remodeling in ICM; yet, the precise composition of the ECM in ICM remains poorly defined. In this study, we employed a sequential protein extraction enabled by the photocleavable surfactant Azo to enrich ECM proteins from left ventricular tissues of patients with end-stage ICM (n=16) and nonfailing donor hearts (n=16). High-resolution mass spectrometry-based quantitative proteomics identified and quantified over 6,000 unique protein groups, including 315 ECM proteins. We discovered significant upregulation of key ECM components, particularly glycoproteins, proteoglycans, collagens, and ECM regulators. Notably, LOXL1, FBLN1, and VCAN were among the most differentially expressed. Functional enrichment analyses revealed enhanced TGFβ signaling, integrin-mediated adhesion, and complement activation in ICM tissues, suggesting a feedback loop driving continued ECM deposition in the end-stage failing heart. Together, our findings provide a comprehensive proteomic landscape of ECM alterations in the end-stage ICM myocardium and identify promising molecular targets for therapeutic intervention.

## Introduction

Ischemic heart disease (IHD) is the most prevalent form of cardiovascular diseases, and remains the leading cause of morbidity and mortality globally.^1–3^ Its incidence continues to rise mainly due to aging population and prevalence of comorbidities such diabetes and obesity, putting an ever-increasing burden on healthcare systems.^1,3^ Ischemic cardiomyopathy (ICM) resulting from chronic coronary artery disease (CAD) and/or acute myocardial infarction (MI) is the most severe clinical manifestation of IHD and the single largest cause of heart failure (HF).^2–4^ With current treatments focused primarily on the management of associated symptoms, there is an urgent need to better understand the molecular changes associated with ischemic HF to enable the development of targeted therapies.

Traditionally, the diagnosis of ICM has been based on a history of myocardial infarction (MI) or objective evidence of coronary artery disease (CAD).^5^ In ICM, cardiomyocyte death resulting from myocardial ischemia leads to remodeling and increased deposition of extracellular matrix (ECM),^6,7^ a process called fibrosis that is primarily driven by myofibroblasts—cardiac fibroblasts activated by growth factors such as TGFβ or increased mechanical stress.^8^ While myocardial fibrosis initially reinforces the ventricular wall—preventing rupture—persistent ECM accumulation ultimately becomes maladaptive, promoting adverse remodeling and compromising cardiac function.^9,10^ The detrimental effects of excessive fibrosis are underscored by its strong association with adverse clinical outcomes, including increased mortality.^11,12^

At its core, the ECM is composed of fibrous proteins such as collagens, fibronectins and laminins; proteoglycans, which consist of core proteins covalently bound to glycosaminoglycan (GAG) chains; secreted factors (e.g., S100 proteins); matrix regulators (e.g., matrix metalloproteinases); and matricellular proteins (e.g., thrombospondins).^6,7,11–15^ Structural components of the cardiac ECM primarily include collagens (types I and III, and to a lesser extent IV, V, and VI), elastin, and fibronectin, whereas nonstructural ECM is largely composed of glycosylated proteins including both proteoglycans and glycoproteins, and glycosaminoglycans such as hyaluronan, heparan sulfate, and chondroitin sulfate.^14,16^ Beyond fibrillar components, cardiac ECM also includes glycoproteins like fibronectin, proteoglycans, GAGs, matricellular proteins, proteases and growth factors. Mass spectrometry-based proteomics has emerged as a powerful tool to define the ECM composition (“matrisome”), enabling identification and quantitation of ECM proteins of healthy and diseased tissues.^17–19^ Proteomics methods have been developed and increasingly applied to investigate cardiac ECM biology and its contributions to disease processes.^13,20–23^ However, the precise compositional difference of the ECM in failing ICM hearts compared to nonfailing donor hearts remain poorly defined.

To address this knowledge gap, we performed proteomic analysis of left ventricular (LV) apex tissues from patients with end-stage failing ICM (n=16) and location-matched samples from nonfailing donor hearts free of cardiovascular disease (n=16). Specifically, we employed sequential protein extraction enabled by photocleavable surfactant 4-hexylphenylazosulfonate (Azo)^24^ to enrich ECM proteins^19,25^ for quantitative mass spectrometry (MS)-based proteomic analysis. Our results revealed striking alterations in the ECM composition of the ICM hearts with significant upregulation of critical ECM components, including glycoproteins, proteoglycans, collagens, and ECM regulators. Notably, LOXL1, FBLN1, and VCAN, which are closely associated with fibrotic remodeling and ECM crosslinking, were among the most differentially expressed. Functional enrichment analyses uncovered activation of TGFβ signaling, integrin-mediated adhesion, and complement pathways, suggesting that that these networks may form a self-sustaining loop driving progressive ECM remodeling. Collectively, our findings provide the first comprehensive snapshot of ECM remodeling in end-stage failing human ICM hearts and identify promising molecular targets for therapeutic intervention to halt or reverse cardiac ECM remodeling and progression to heart failure.

## Methods

### Chemicals and reagents

Trypsin Gold, Mass Spectrometry Grade was purchased from Promega (Madison, WI, USA; catalog no. V5280). The photocleavable surfactant Azo was purchased from MilliporeSigma (Burlington, MA, USA; catalog no. 919233). Solutions were prepared in HPLC-grade water from Fisher Scientific (Fair Lawn, NJ, USA). Amicon® Ultra Centrifugal Filter, 10 kDa MWCO filters were purchased from MilliporeSigma (catalog no. UFC5010). All other reagents and buffers were purchased from Fisher Scientific or MilliporeSigma unless otherwise noted.

### Human Cardiac Tissue Collection

Available clinical data including age, gender, cause of death, and medical history are listed for the heart tissues used in this study in **Table S1**. Tissue from the apex of the LV of failing ICM and nonfailing donor hearts (n=16 each) was used for the experiments described in this study. Tissue from failing ICM hearts was obtained from individuals undergoing left ventricular assist device (LVAD) implantation surgery at the University of Michigan Medical Center. ICM was diagnosed based on known CAD severity that would account for HF. Nonfailing donor hearts (n=16) were obtained from the University of Wisconsin-Madison Organ and Tissue Donation Program. Donor hearts were stored in cardioplegic solution following removal then dissected and flash frozen in liquid nitrogen for storage at -80 °C. The procedures for the collection of human failing ICM and nonfailing donor heart tissues were approved by the Institutional Review Boards of the University of Wisconsin-Madison and the University of Michigan, respectively.

### Protein Extraction

To extract proteins, approximately 15 mg of myocardial tissue from failing ICM and nonfailing donor hearts were cut and pulverized under liquid nitrogen using a CellCrusher kit (Fisher Scientific; catalog no. NC1824866). Cryopulverized tissue was resuspended in 1X Dulbecco’s Phosphate-Buffered Saline (DPBS; MilliporeSigma; catalog no. D1283) supplemented with Halt Protease Inhibitor Cocktail (Fisher Scientific; catalog no. 87786) diluted to a final concentration of 1X by vortexing for 10 sec and then centrifuged at 21,100 × *g* for 5 min. The resulting supernatants (wash) were removed and discarded. To achieve deeper coverage of the cardiac proteome by mass spectrometry-based proteomics, sequential extraction of proteins was performed with the remaining pellets. Pellets were homogenized in 20 volumes (approx. 300 uL) of LiCl extraction buffer [3 M LiCl, 1 mM TCEP, 10 mM EDTA, and 1X Halt Protease Inhibitor Cocktail (Fisher Scientific)] using a handheld Teflon homogenizer (Bel-Art, Wayne, NJ, USA). Homogenates were vortexed and centrifuged as before and the resulting supernatants (LiCl extracts) were desalted using Ultra Centrifugal Filter, 10 kDa MWCO filters (MilliporeSigma). The pellets remaining after LiCl extraction were re-homogenized in 20 vol. (approx. 300 uL) of Azo extraction buffer [0.4% Azo (MilliporeSigma), 25 mM ammonium bicarbonate, 1 mM TCEP, 10 mM EDTA, and 1X Halt Protease Inhibitor Cocktail (Thermo Scientific)] using a handheld Teflon homogenizer (Bel-Art). Homogenates were subsequently sonicated first via probe sonication (Fisher Scientific; 20% amplitude, 3 cycles, 3 sec per cycle) followed by incubation in an ultrasonication bath (Fisher Scientific) for 30 min, followed by incubation in an Eppendorf Thermomixer R (Hamburg, Germany) at 80 °C for 15 min. This extract was centrifuged at 21,100 × *g* for for 20 min then the supernatant was saved and referred to as the Azo extract.

### Protein Digestion and Desalting

All extracts (n=16 control/LiCl 1; n=16 control/Azo; n=16 ICM/LiCl 1; n=16 ICM/Azo) were normalized to approximately 75 µg of total protein via Bradford protein assay and diluted to the same final concentration (1 mg/mL). LiCl extracts were diluted with 25 mM ammonium bicarbonate, and Azo extracts in the remaining 0.4% Azo extraction buffer. For reduction and alkylation of disulfide bonds, samples were treated simultaneously with 25 mM TCEP and 50 mM chloroacetamide and incubated at 37 °C and 600 rpm in a Thermomixer R (Eppendorf) for 30 min, after which they were treated with 1 M ammonium bicarbonate to adjust pH to the active range for digestion (∼pH 8.5). Digestion was performed by treating samples with a 50:1 (w/w) protein:trypsin ratio and incubating them overnight in a Thermomixer R (Eppendorf) at 37 °C and 600 rpm. Digestion was quenched by addition of formic acid to an approximate final concentration of 0.25% (pH 3-4). Extracts containing Azo were irradiated in front of a UV lamp to degrade surfactant, then centrifuged for 30 min. All extracts were then desalted by Pierce C18 tips (ThermoFisher Scientific; catalog no. 87784) to remove degradation products and other salts, dried, and reconstituted in 0.1% formic acid in water. The peptide content of each sample was assessed using a NanoDrop™ One/One^C^ Microvolume UV-Vis Spectrophotometer (Thermo Scientific).

### Data Acquisition

A Bruker timsTOF Pro trapped ion mobility Q-TOF instrument (Billerica, MA, USA) fitted with a captive-spray nano-electrospray ionization source and coupled to a nanoElute nanoflow LC system (Bruker) was used for all analyses. For each sample, a total of 200 ng of peptides was injected onto an Aurora Ultimate C18 column (25 cm length, 75 μm inner diameter, 1.7 μm particle size, 120 Å pore size; IonOpticks, Collingwood, Australia). Injections were normalized to achieve a total ion chromatogram (TIC) intensity of approximately 3e7 for each sample. Separations were carried out at a flow rate of 0.4 µL/min and 55 °C using a linear gradient increasing from 0 to 17% mobile phase B (0.1% formic acid in acetonitrile) (mobile phase A: 0.1% formic acid in water) over 60 min; 17% to 25% from 60 to 90 min; 25% to 37% B from 90 to 100 min; 37% to 85% B from 100 min to 110 min; and a 10 min hold at 85% B before washing and returning to low organic conditions. Mass and tandem mass spectra were obtained in diaPASEF mode, with 32 fragmentation windows from 400-1200 m/z (25 m/z width) and 0.6 to 1.6 1/K0 (0.3 1/K0 width).

### Database Searching and Quantitation

Raw MS data were searched in a library-free manner using DIA-Neural Network^26^ version 1.8.1 with the following parameters: 1% FDR, Library-free search enabled, Minimum fragment m/z: 200, Maximum fragment m/z: 2000, N-terminal methionine cleavage enabled, Enzyme: Trypsin, Maximum missed cleavages: 1, Minimum peptide length: 7, maximum peptide length: 40, Minimum precursor m/z: 400, Maximum precursor m/z: 1400, Minimum precursor charge: 1, Maximum precursor charge: 5, cysteine carbamidomethylation enabled, MS1/MS2 mass accuracy: 10 ppm, Quantification strategy: Robust LC (High Precision), Neural network classifier: Double-pass mode. All other parameters were left as defaults. Data were searched against a FASTA file containing 20,379 canonical human protein sequences downloaded from UniprotKB. A combined list of all identified protein groups and corresponding peptides are listed in **Tables S2 and S3**, respectively.

Downstream differential expression analysis was performed as previously described.^27^ Quantified proteins were filtered for validity using the “DAPAR”^28^ package for R to include all proteins quantified in 8 of 16 runs in at least one sample group. After median normalization, imputation of missing values was performed via the ssla algorithm (values partially observed in some replicates of a condition) or set to the quantile below which 2.5% of all observations fall (values that were missing entirely within a condition). Identifications were further filtered to remove any proteins which had imputed values for more than 50% of their quantities in each extract. The “DEP” R package was used to perform a limma test between all specified contrasts,^29,30^ and the “IHW”^31^ R package was used to adjust all p-values, using the number of quantified peptides per protein as a covariate. The criteria for significance were 1) an adjusted p-value less than 0.05 and 2) a log2 fold change greater than or equal to 0.6 (approximately equivalent to a 1.5-fold change in the normal scale).

### Gene Ontology (GO) and Pathway Analysis

For GO analysis, proteins differentially expressed in ICM relative to nonfailing donor myocardium were searched in STRING^32^ and the functional enrichment categories for biological processes and molecular functions were exported into RStudio. GO terms were filtered to a 1% false discovery rate and plotted using “ggplot2” and “ggpubr” packages.^33^ Enrichment analysis using active subnetworks was further performed using the package “pathfinder”^34^ using the Kyoto Encyclopedia of Genes and Genomes (KEGG) as the gene set and a p-value threshold of 5%. Results were plotted similarly to the GO analyses. Protein-protein interaction networks from STRING were visualized using Cytoscape.^35^ ECM proteins and categorical information were retrieved from MatrisomeDB.^36^

## Results

### Global Proteomics Analysis of Failing ICM and Nonfailing Donor Tissues

To identify differences in the composition of the ECM in failing ICM versus nonfailing donor myocardium, we optimized our cardiac tissue extraction protocol to maximize solubilization of ECM proteins. Our group had previously demonstrated the efficacy of using the photocleavable surfactant, Azo, alone and in combination with a decellularization step using a highly concentrated salt such as NaCl in a sequential extraction.^19,25^ However, this approach had not been applied to fibrotic, dense muscle tissues such as the human cardiac tissue. Prior studies have reported the improved performance of LiCl over NaCl in the extraction of proteins from muscle tissues,^37^ and our group has successfully used LiCl to extract myofibrillar proteins from human heart tissue.^38^ Thus, we employed sequential extractions first with LiCl followed by Azo-containing buffer to effectively decellularize tissue and solubilize ECM proteins.

To gain insight into differences in the composition of the ECM in end-stage ICM, this optimized sequential protein extraction protocol was employed to decellularize tissue and solubilize ECM proteins from the apex of the LV myocardium of failing ICM (n=16) and nonfailing donor (n=16) hearts (**Figure 1A)**. The LiCl and Azo extracts (64 total samples; n=32 ICM, n=32 control) were subjected to analysis by MS-based proteomics. The total ion chromatograms from each extract across cohorts and normalized intensities following searching demonstrated consistency between biological replicates (**Figure S1 and S2**).

**Figure 1.**
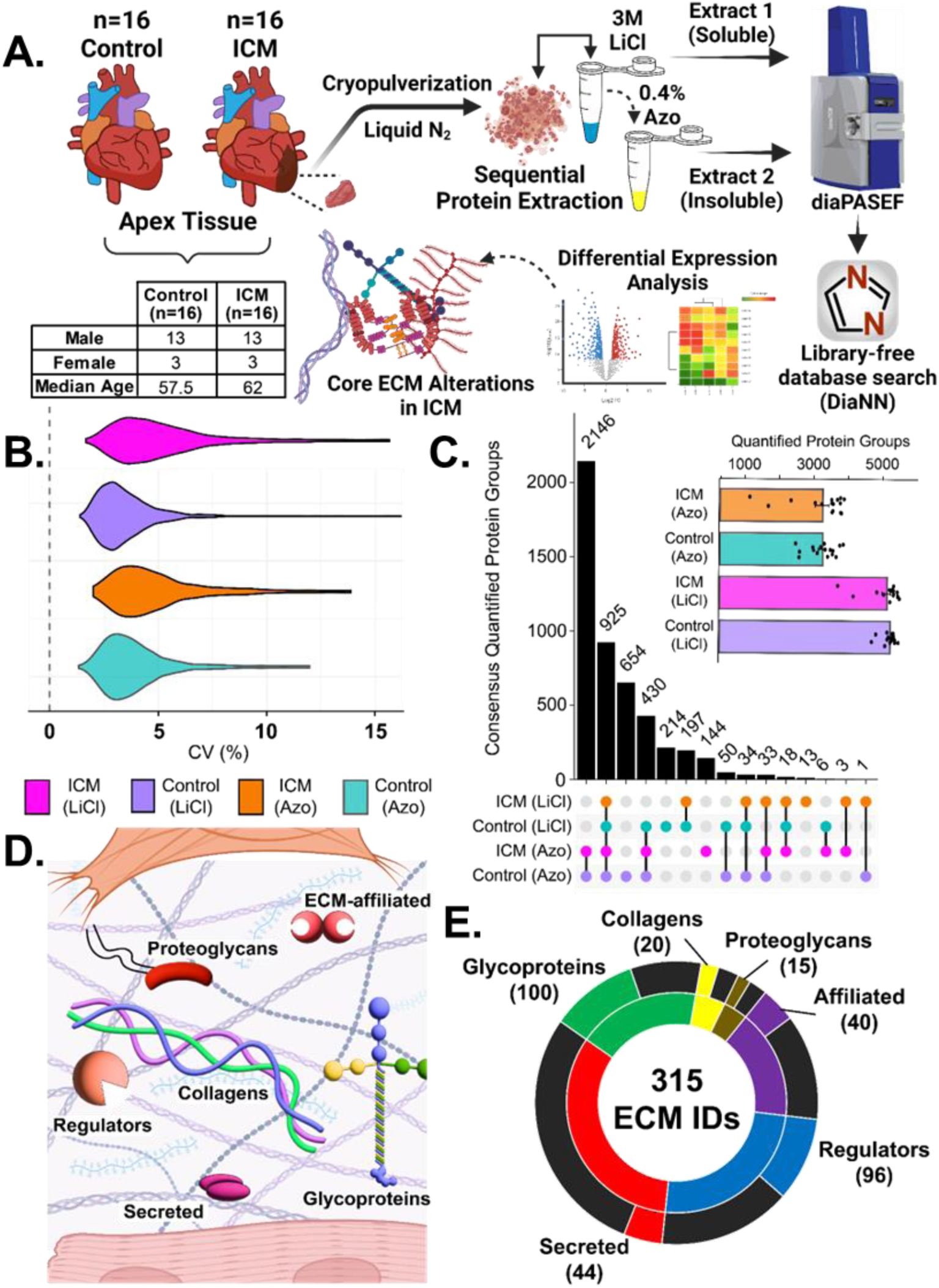
Azo-enabled ECM proteomics method yields reproducible coverage of the matrisome (the collection of proteins in the ECM) in failing ICM and nonfailing donor heart tissues. **A)** Schematic representation of the workflow for MS-based proteomic analysis of protein extracts from human ICM and donor patient myocardial tissue. Soluble proteins are depleted in the first LiCl extraction (LiCl) followed by solubilization of “insoluble” ECM proteins by extraction with buffer containing the photocleavable surfactant, Azo. **B)** CV of protein intensities across extracts from failing ICM and nonfailing donor tissues. The median CV of protein intensities was below 5% for extracts from both groups, indicating high quantitative reproducibility. **C)** Upset plot and bar graph (inset) showing >6000 unique protein identifications overall, as well as the degree of overlap between groups. **D and E)** Simplified depiction of major categories of ECM proteins (per MatrisomeDB^36^) **(D)** and breakdown of the number of proteins in each category identified in this study **(E)**. The inner circle shows the total number of proteins in each category while the outer circle indicates those that were identified herein. Black segments correspond to proteins present in MatrisomeDB^36^ that were not identified in human ICM and donor myocardial tissue.

For each of the extracts and disease conditions, quantitative reproducibility was assessed by plotting the coefficients of variation (CV) for the protein-level normalized label-free quantified (LFQ) intensities (**Figure 1B**). All groups showed median CV values of <5%. In addition, normalized LFQ intensities were plotted for each replicate against the others within each of the four sample groups and showed an average Pearson correlation coefficient of r = 0.948. For ICM samples, the average r values for the LiCl and Azo extracts were 0.936 and 0.93, respectively, while for nonfailing controls they were 0.966 and 0.96 (**Figure S3**).

Following database searching of raw mass and tandem mass spectra, data were filtered to retain only genes with 8 or more valid quantified values in at least one of the conditions, resulting in the identification of a total of 6,125 unique protein groups across both extracts. After a second filtration step wherein proteins with more than 50% imputed values were removed, 6,118 unique protein groups remained. Overall, more protein groups were quantified in the LiCl extract than the Azo extract, though Azo exhibited a greater degree of overlap between ICM and control donor tissues suggesting better consistency in detecting shared proteins (**Figure 1C**). Notably, when comparing the protein intensities across all extracts, we found that major fibrillar proteins, such as type 1 collagen, fibronectin, and elastin, were significantly more abundant in Azo extracts, showing our method effectively enriched insoluble ECM proteins (**Figure S4**). To assess the effectiveness of ECM extraction, the total number of ECM proteins in all extracts were identified according to information in MatrisomeDB^36^ and plotted according to their category. These categories consist of collagens, proteoglycans, glycoproteins, ECM regulators, secreted proteins, and ECM-affiliated proteins (**Figure 1D**). These results demonstrated that our method was effective at extracting all categories of ECM proteins, with particularly high coverage of glycoproteins in which our dataset captured ∼55% of those in the database (**Figure 1E**).

### Alterations in Failing Human ICM Myocardium Revealed by Global Proteomics

Principal component analysis (PCA) of the Azo extract showed clear separation between failing ICM and nonfailing donor samples along PC1, which account for 33.34% of the total variance (**Figure 2A**). Similar clustering by condition was observed in the PCA results of the LiCl extract, though the separation explained a smaller proportion of the variance (**Figure S5**). In general, ICM samples had less cohesion in PCA clustering which alongside lower Pearson correlation coefficients can be explained by the relatively higher degree of heterogeneity in the ICM tissues compared to controls. This can likely be explained by differences in factors such as prior MI status, treatments, and comorbid conditions like diabetes (**Table S1**). Differential expression analysis was performed by a limma test, yielding 313 differentially expressed protein groups in the Azo extract (of 2,489 quantified protein groups) and 826 differentially expressed protein groups in LiCl. These were sorted in descending order of adjusted p-value, then the top 300 results were plotted in a hierarchical heatmap and clustered by k-means, revealing a distinct pattern in protein abundance in failing ICM compared to nonfailing donor myocardium (**Figure 2B**).

**Figure 2.**
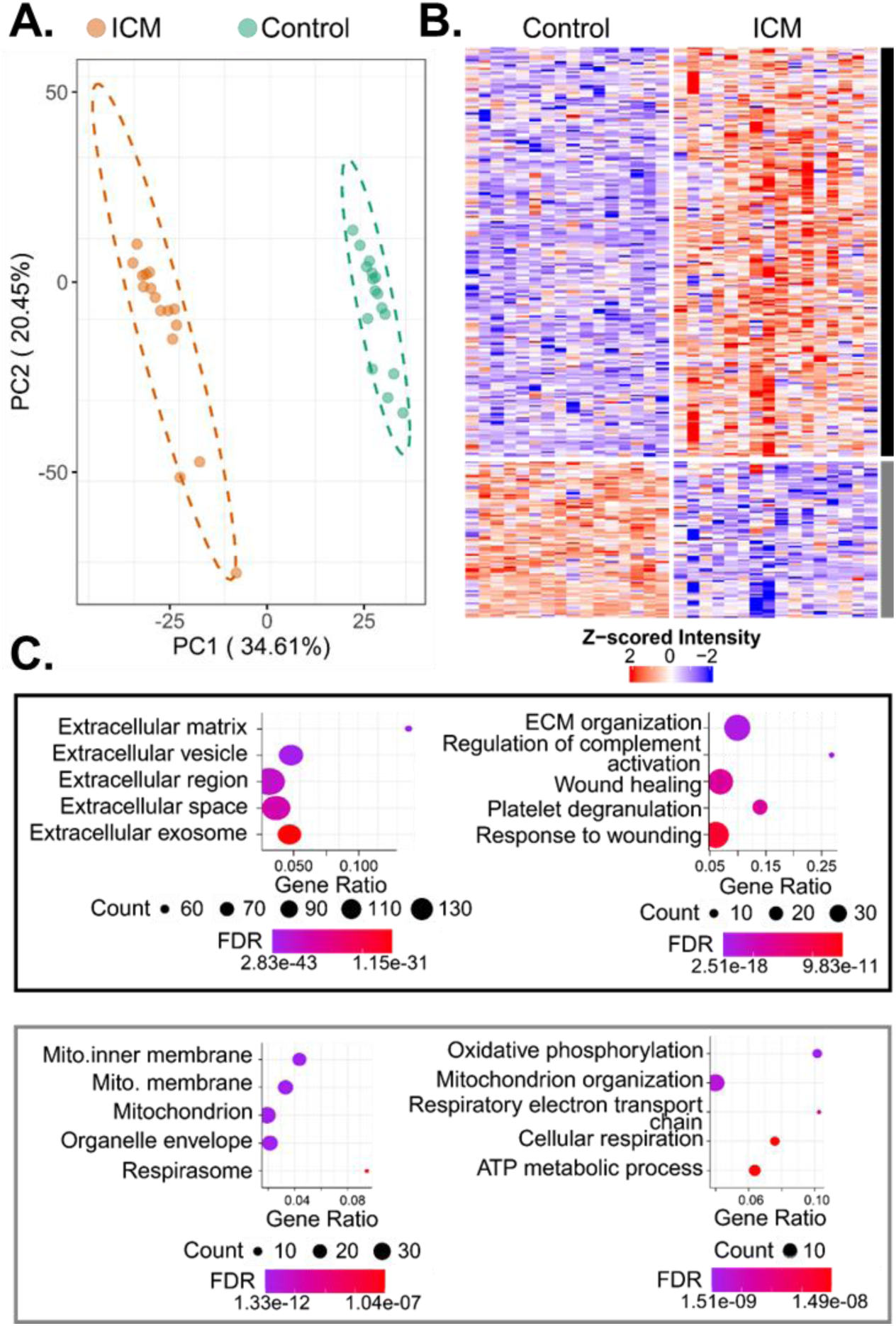
Global proteomics results from ECM-enriched extract reveals alterations in failing ICM versus nonfailing donor myocardium. **A)** PCA shows clustering and clear separation of failing ICM and nonfailing donor group replicates. **B)** Heatmap of Z-scored protein intensities (top 300 most significant differentially expressed) shows clusters (k-means) of proteins up- and down-regulated in failing ICM versus nonfailing control myocardium. **C)** Bubble plots showing GO terms enriched among protein clusters up- (black) and down-regulated (grey) in end-stage failing ICM compared to nonfailing control. Enriched cellular component terms (left) highlight up- and down-regulation of ECM and mitochondrial proteins, respectively, in failing ICM compared to nonfailing control myocardium. Similarly, biological process terms (right) enriched among proteins up-regulated in the failing ICM myocardium pertained to ECM organization and complement activation while those enriched among down-regulated proteins were concerned with metabolic processes.

Differentially expressed proteins broadly clustered into two groups: those that were up- and those that were down-regulated in failing ICM samples relative to nonfailing controls. GO analysis was performed for differentially expressed proteins, and the GO terms, associated proteins, enrichment strength, and false discovery rate (FDR) are reported in **Table S4**. GO analysis of the 225 proteins up-regulated in failing ICM versus nonfailing donor myocardium showed unambiguous association with the ECM (GO:0031012; FDR= 2.83E-43) and its functions, including ECM organization (GO:0030198; FDR= 2.51E-18), wound healing (GO:0042060; FDR= 4.59E-12), and regulation of complement activation (GO:0030449; FDR= 1.68E-10). Other enriched terms were related to blood coagulation (GO:0007596; 9.83E-11), platelet degranulation (GO:0002576; FDR=3.9E-11), proteolysis (negative regulation of endopeptidase activity; GO:0010951; 6.92E-10), and cell adhesion (GO:0007155; FDR=1.47E-05). The 86 down-regulated proteins were associated with the mitochondrion (GO:0005739; FDR=5.48E-11) and oxidative phosphorylation (GO:0006119; FDR=1.51E-09), underscoring the established link between mitochondrial dysfunction and HF^3,39^ (**Figure 2C and 2D**). GO analysis from the LiCl extract exhibited nearly identical trends, with more terms related to cytosolic proteins attributable to the role of this step in decellularization of tissues (**Figure S6**).

### Cardiac ECM remodeling in end-stage HF from ICM

Global proteomics results showed that many of the detected ECM proteins (68 out of the total 315 ECM proteins detected) were up-regulated in failing ICM compared to nonfailing donor heart tissue. We contextualized these findings using information in MatrisomeDB^36^ to outline categories by which to observe matrisome changes in ICM. In total, 315 ECM proteins were identified from the raw data search, which reduced to 286 after all filtering steps, consisting of 89 glycoproteins (46.6% of database), 18 collagens (40%), 14 proteoglycans (38.9%), 37 ECM-affiliated proteins (22.9%), 87 ECM regulators (35%), and 41 secreted proteins (12.4%). To visualize the categorical ECM differences in ICM compared to control hearts, we plotted the quantile function (0.95% confidence interval) of the log2 fold changes of differentially expressed ECM proteins across each matrisome category (**Figure 3A**). Categories that were considered significantly up-regulated in failing ICM versus nonfailing control tissues had a mean fold change (± the confidence interval) greater than 0, indicating that glycoproteins (p-value=5.23E-11), ECM regulators (p-value=2.29E-05), proteoglycans (p-value=1.62E-05), and collagens (p-value=4.02E-05) showed significantly higher abundance in Azo of the ICM samples categorically. The same categories were significant in LiCl with the exceptions of ECM-affiliated proteins and collagens, the latter of which we can assume is attributable to increased collagen solubilization by the Azo surfactant. **Table 1** summarizes the p-values and log2 fold changes for several categories of ECM proteins significantly up-regulated in the failing ICM versus nonfailing control myocardium based on the Azo data. The individual proteins from the table are plotted in categories as box plots showing individual replicates in **Figures S7-S10**.

**Figure 3.**
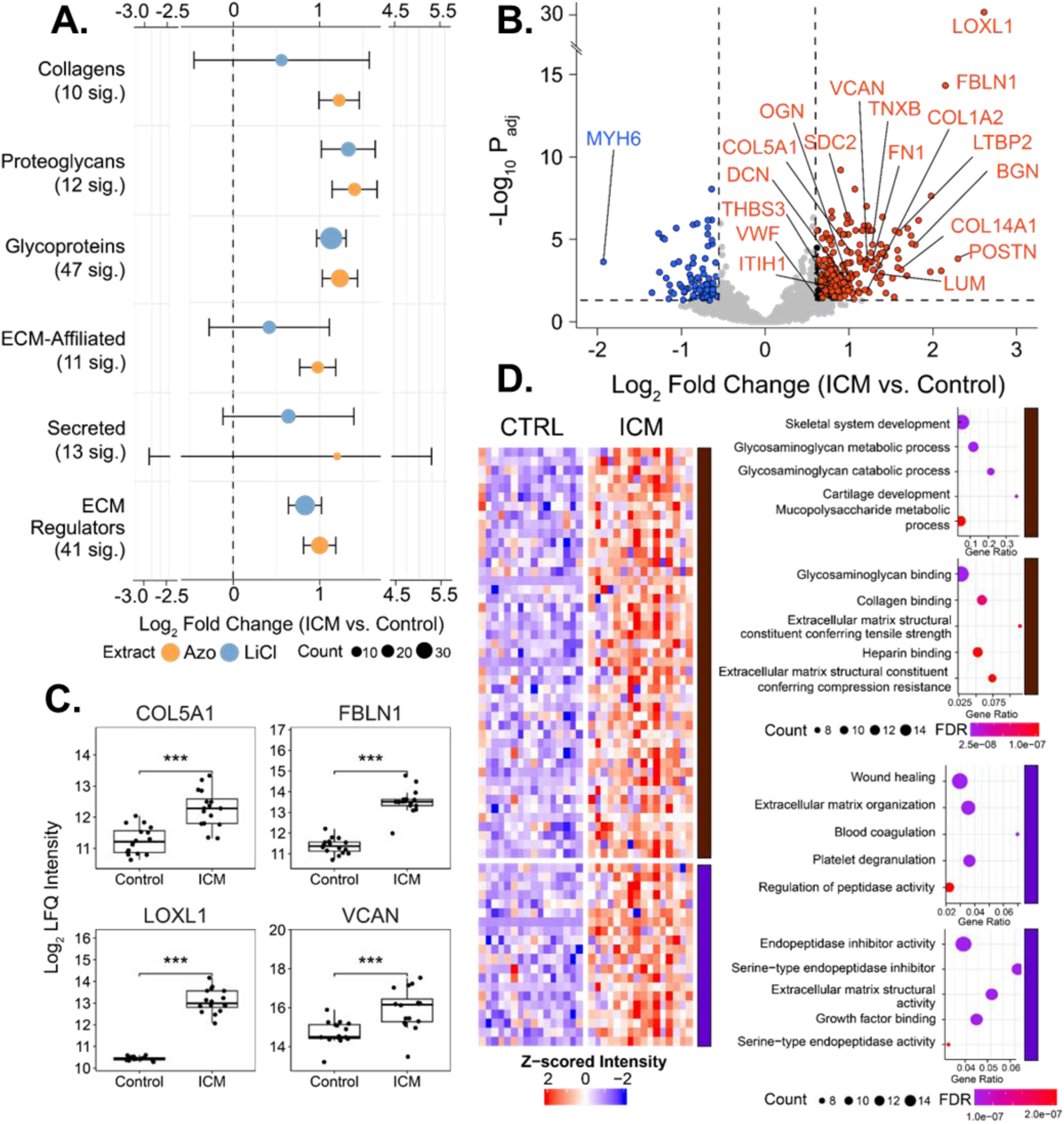
ECM remodeling in end-stage HF from ICM. **A)** Dot plot showing categorical log2 fold change in the intensities of matrisome proteins. Mean and standard deviation of log2 fold changes for all ECM proteins in each extract and category were plotted. Overall, core matrisome proteins (collagens, proteoglycans, and glycoproteins), as well as ECM regulators, were significantly upregulated in failing ICM LV tissue. **B)** Volcano plot showing select proteins up- and down-regulated in failing ICM versus nonfailing donor myocardium. **C)** Core collagens, ECM regulators, glycoproteins and proteoglycans show increased expression in ICM. **D)** Clustered heatmap and corresponding bubble plots showing the results of GO analysis using the list of ECM proteins up-regulated in failing ICM myocardial tissue. In the first cluster, biological processes (top) relate to carbohydrate metabolism with molecular functions (bottom) of binding and matrix structural properties. The second cluster pertains more to matrix regulation by growth factors and peptidases.

**Table 1.**
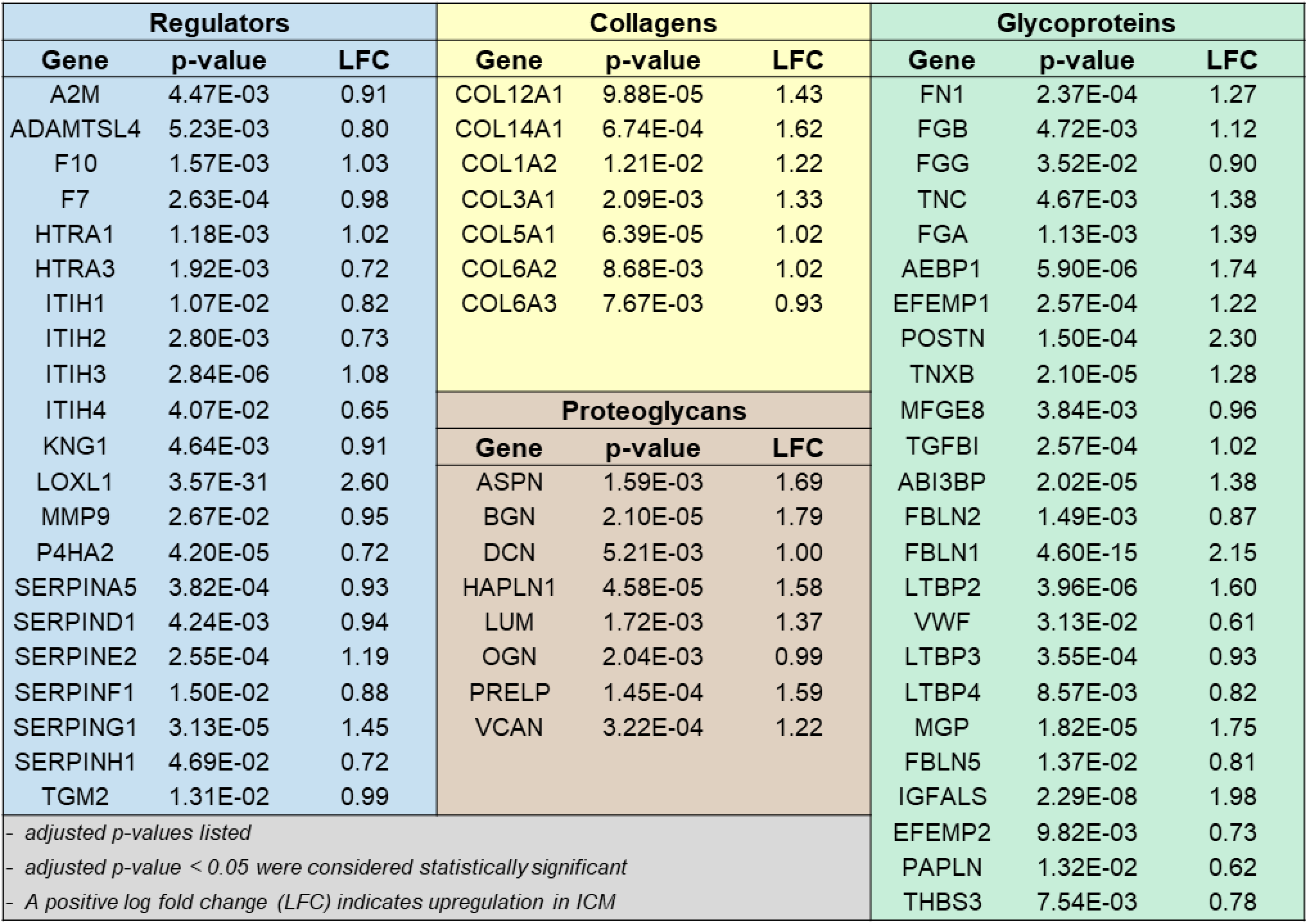
Proteins significantly up-regulated in the Azo extract from failing ICM versus nonfailing donor hearts.

| Regulators |  |  | Collagens |  |  | Glycoproteins |  |  |
| --- | --- | --- | --- | --- | --- | --- | --- | --- |
| Gene | p-value | LFC | Gene | p-value | LFC | Gene | p-value | LFC |
| A2M | 4.47E-03 | 0.91 | COL12A1 | 9.88E-05 | 1.43 | FN1 | 2.37E-04 | 1.27 |
| ADAMTSL4 | 5.23E-03 | 0.80 | COL14A1 | 6.74E-04 | 1.62 | FGB | 4.72E-03 | 1.12 |
| F10 | 1.57E-03 | 1.03 | COL1A2 | 1.21E-02 | 1.22 | FGG | 3.52E-02 | 0.90 |
| F7 | 2.63E-04 | 0.98 | COL3A1 | 2.09E-03 | 1.33 | TNC | 4.67E-03 | 1.38 |
| HTRA1 | 1.18E-03 | 1.02 | COL5A1 | 6.39E-05 | 1.02 | FGA | 1.13E-03 | 1.39 |
| HTRA3 | 1.92E-03 | 0.72 | COL6A2 | 8.68E-03 | 1.02 | AEBP1 | 5.90E-06 | 1.74 |
| ITIH1 | 1.07E-02 | 0.82 | COL6A3 | 7.67E-03 | 0.93 | EFEMP1 | 2.57E-04 | 1.22 |
| ITIH2 | 2.80E-03 | 0.73 | Proteoglycans |  |  | POSTN | 1.50E-04 | 2.30 |
| ITIH3 | 2.84E-06 | 1.08 |  |  |  | TNXB | 2.10E-05 | 1.28 |
| ITIH4 | 4.07E-02 | 0.65 |  |  |  | MFGE8 | 3.84E-03 | 0.96 |
| KNG1 | 4.64E-03 | 0.91 |  |  |  | TGFBI | 2.57E-04 | 1.02 |
| LOXL1 | 3.57E-31 | 2.60 |  |  |  | ABI3BP | 2.02E-05 | 1.38 |
| MMP9 | 2.67E-02 | 0.95 |  |  |  | FBLN2 | 1.49E-03 | 0.87 |
| P4HA2 | 4.20E-05 | 0.72 |  |  |  | FBLN1 | 4.60E-15 | 2.15 |
| SERPINA5 | 3.82E-04 | 0.93 |  |  |  | LTBP2 | 3.96E-06 | 1.60 |
| SERPIND1 | 4.24E-03 | 0.94 |  |  |  | VWF | 3.13E-02 | 0.61 |
| SERPINE2 | 2.55E-04 | 1.19 |  |  |  | LTBP3 | 3.55E-04 | 0.93 |
| SERPINF1 | 1.50E-02 | 0.88 | ASPN | 1.59E-03 | 1.69 | LTBP4 | 8.57E-03 | 0.82 |
| SERPING1 | 3.13E-05 | 1.45 | BGN | 2.10E-05 | 1.79 | MGP | 1.82E-05 | 1.75 |
| SERPINH1 | 4.69E-02 | 0.72 | DCN | 5.21E-03 | 1.00 | FBLN5 | 1.37E-02 | 0.81 |
| TGM2 | 1.31E-02 | 0.99 | HAPLN1 | 4.58E-05 | 1.58 | IGFALS | 2.29E-08 | 1.98 |
| - adjusted p-values listed<br>- adjusted p-value < 0.05 were considered statistically significant<br>- A positive log fold change (LFC) indicates upregulation in ICM |  |  | LUM | 1.72E-03 | 1.37 | EFEMP2 | 9.82E-03 | 0.73 |
|  |  |  | OGN | 2.04E-03 | 0.99 | PAPLN | 1.32E-02 | 0.62 |
|  |  |  | PRELP | 1.45E-04 | 1.59 | THBS3 | 7.54E-03 | 0.78 |
|  |  |  | VCAN | 3.22E-04 | 1.22 |  |  |  |

Plotting the log2 fold changes and adjusted p-values (with thresholds of 0.6 and 0.05, respectively) reveals that the collagen and elastin crosslinking enzyme LOXL1 (lysl oxidase-like protein 1) is the most highly up-regulated protein in the failing ICM, with the lowest p-value (3.57E-31) and second-largest log2 fold change (2.60) (**Figure 3B-C**). In other disease models including idiopathic pulmonary fibrosis and liver cirrhosis, LOXL1 was shown to be stimulated by pro-fibrotic TGFβ to increase crosslinking and that its knockdown protects from fibrosis and reduces expression of pro-fibrotic metalloproteases and collagens.^40^ The next most significantly up-regulated protein based on p-value was the adhesive ECM glycoprotein fibulin-1 (FBLN1; p-value=4.6E-15), which also had the fourth-highest log2 fold change overall in the Azo extract (**Figure 3B-C**). During cardiac development, FBLN1 promotes ADAMTS-1 cleavage of chondroitin sulfate proteoglycan VCAN (versican).^41^ VCAN accumulation has been previously reported in failing hearts and shown to be reduced upon treatment with β-blockers regulated by ADAMTS family proteins. ^21^ Consistent with this, we observed significantly higher VCAN expression in the failing ICM compared to nonfailing donor tissue (p-value=3.22E-04) (**Figure 3B-C**). These findings suggest that the marked increase in FBLN1 observed in failing ICM may serve as a protective mechanism, potentially compensating for elevated VCAN expression by stimulating ADAMTS-mediated proteolysis.

To further probe the pathways underlying these ECM differences in ICM, we performed hierarchical clustering of the ECM proteins significantly up-regulated in failing ICM compared to nonfailing donor myocardium, again with two k-means clustering (**Figure 3D**). In Cluster 1 we observed many terms related to glycosaminoglycans (GAGs), including GAG metabolism (GO:0030203; FDR=6.13E-08), catabolism (GO:0006027; FDR=1.05E-07) and sulfur compound metabolic process (GO:0044273; FDR=1.44E-06) relating to GAG sulfation. Of note, there were several terms related to cartilage development (GO:0051216; FDR=1.05E-07). Evidence shows similar hierarchical organization and regulation of ECM in both heart and cartilaginous connective tissues, including regulatory pathways involving ACAN (aggrecan) and tenascins (TNC and TNXB), which we found were significantly up-regulated in ICM tissues.^42,43^ The GO functional enrichment analysis returned terms primarily related to binding – collagen binding (GO:0005518; FDR=1.26E-09), GAG binding (GO:0005539; FDR=1.33E-12), integrin binding (GO:0005178; FDR=1.40E-04), ECM binding (GO:0050840; FDR=1.30E-03), and calcium ion binding (GO:0005509; FDR=2.32E-06) – though also had terms related to specific matrix structural elements. ECM structural constituents conferring tensile strength (i.e. resistance to stretching) were all collagens (COL1A2, COL3A1, COL6A2, COL6A3, COL12A1, COL14A1; GO:0030020; FDR=2.93E-08), while those conferring compression resistance were proteoglycans (DCAN, VCAN, LUM, BGN, PRELP; GO:0030021; FDR=1.28E-07). Cluster 2 terms overlapped greatly with the generally up-regulated protein groups, with major categories such as coagulation (e.g. negative regulation of blood coagulation; GO:0030195; FDR= 4.66E-06) and proteolysis (GO:0052547; FDR=2.86E-07). Functional terms make it clear these trends relate to the SERPIN family of serine protease inhibitors (GO:0004867; FDR=8.18E-06) and matrix metalloproteinases, MMP9 in particular. Growth factor signaling also appears as an enriched term, which is consistent with the role of TGFβ in promoting ECM remodeling and fibrosis in ICM.^44,45^

### Pathway Analysis Revealed TGFβ Signaling Drives ECM Remodeling in End-Stage ICM

We next performed functional annotation clustering using DAVID^46^ (Database for Annotation, Visualization and Integrated Discovery) to group similar annotations and clarify broader trends in the differentially expressed proteins in ICM tissues (**Table S4**). Clusters were ranked by enrichment score, a geometric mean of p-values for terms within a cluster and using the differentially expressed proteins in the Azo extract we observed 20 clusters wherein the top term’s Benjamini-Hochberg adjusted p-value was significant (<0.05). The top two clusters (enrichment scores=22.77 and 16.19) unsurprisingly contained KEGG and GO terms relating to ECM components, though the second more generically related to ECM components and the first had annotations of secreted proteins and glycoproteins. The next highest clusters were related to blood coagulation (enrichment score=6.13), cell adhesion (5.12), calcium-binding epidermal growth factor (EGF) domains (3.49), and SERPINs (3.48). Clusters 9 and 10 reflected complement pathways/innate immunity (2.97) and immunoglobulin binding (2.92), and Cluster 12 reflected SLRPs, such as DCN, LUM, BGN, and PRELP (2.58).

To provide the context of protein-protein interactions into the pathway analyses, we used the R package “pathfinder”^34^ to perform an active subnetwork search in a protein interaction network, then perform pathway enrichment using these networks. A p-value threshold of 0.05 was used for significance, and active subnetwork search is iterated ten times, with the lowest p-value amongst iterations used to determine the overall significance of the term. Using KEGG annotations, 51 terms were significantly enriched from Azo using this approach and the terms with the largest fold changes were complement and coagulation cascades (lowest p-value=1.06E-21), oxidative phosphorylation (p-value=3.04E-26), cardiac muscle contraction (p-value=3.87E-10), and ECM-receptor interaction (p-value=3.28E-11) (**Figure 4A**).

**Figure 4.**
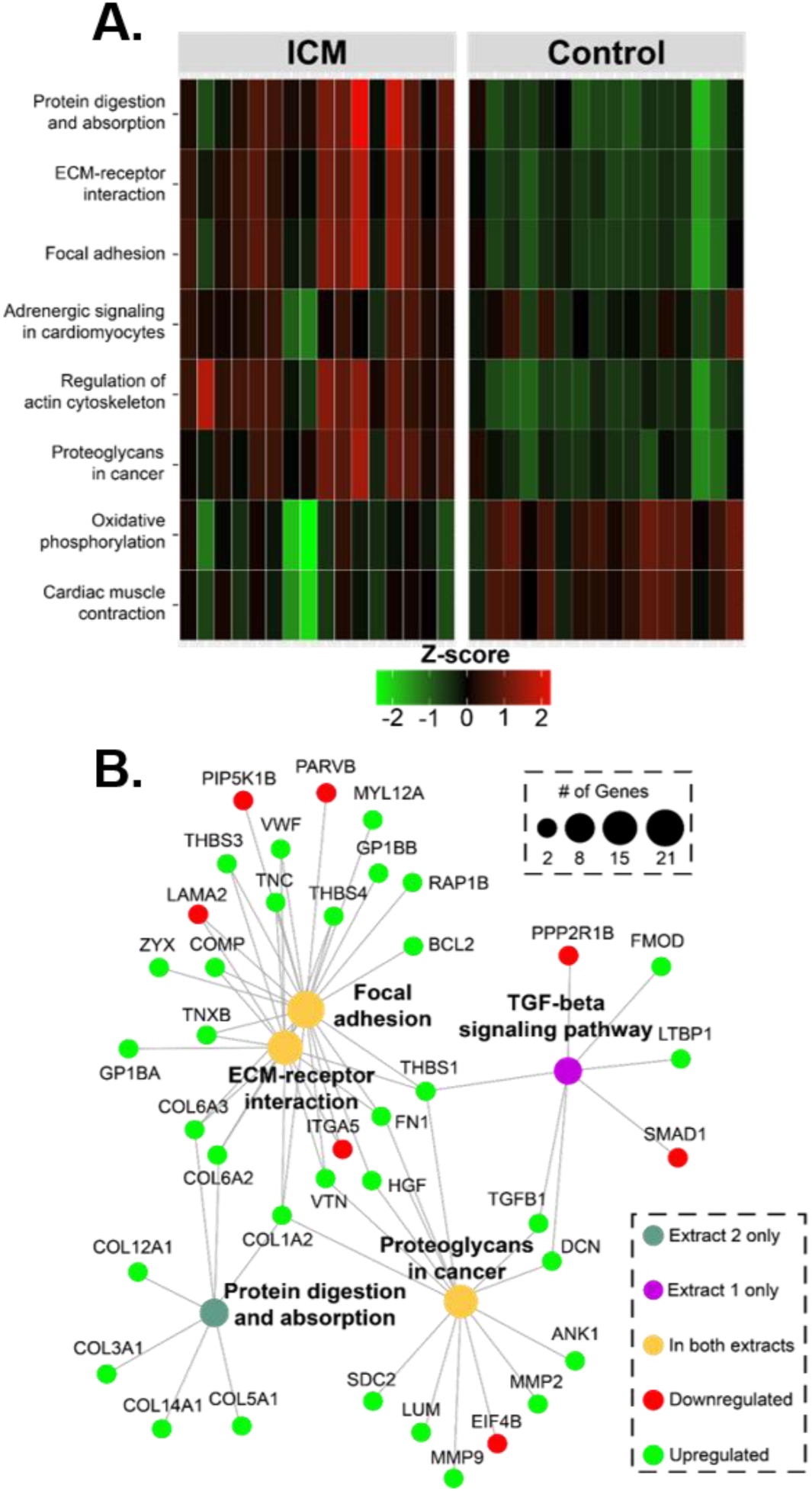
Pathway analysis implicates TGFβ signaling network in altered ECM composition in the myocardium of individuals with end-stage failing ICM. **A)** Heatmap of agglomerated Z-scores for KEGG pathway terms across samples show up-regulation of pathways related to focal adhesions and ECM-receptor interactions. In addition, these changes are accompanied by increased regulation of the cytoskeleton and decreased expression of metabolic and contractile proteins. Term changes are visualized for the ECM-enriched extract. **B)** Term-protein network generated using input data from both extracts demonstrates connections between pathways via specific proteins. In general, adhesive glycoproteins (thrombospondins, tenascins, fibronectin, vitronectin) and proteoglycans (decorin), connect receptor processes to focal adhesion and regulation by protein digestion and TGFβ pathways.

From GO terms (using all terms from cellular component, biological process, and molecular function), 92 entries were deemed significant. Significant GO terms were largely in agreement with the previous GO results, though the fold enrichment provided for each term provided new context for particularly large changes, including the 61-fold enrichment for ECM structural constituents conferring compression resistance (p-value=2.38E-2), a category containing mainly small leucine-rich proteoglycans. Notably, integrin activation (p-value=2.72E-5) and integrin-mediated signaling pathways (p-value=3.39E-05) were also significant, displaying a 24-fold and a 10-fold enrichment respectively. Hierarchical clustering of the enriched terms linked ECM organization to integrin binding (**Figure S11**), and plotting individual proteins showed FN1, APOE, and EFEMP2 linked those terms as well as endoplasmic reticulum lumen, heparin binding, and peptide cross-linking (**Figure S12**).

Reactome searches yielded 134 significant terms of which the majority with the lowest p-values were related to inflammation, hemostasis, and dysregulated electron transport. Insulin-like growth factor binding was previously shown as a significant term from GO analyses, and a similar Reactome annotation, “Regulation of Insulin-like Growth Factor (IGF) transport and uptake by Insulin-like Growth Factor Binding Proteins (IGFBPs)”, provided a more specific description of the pathway with an expanded list of relevant proteins. Hierarchical clustering showed multiple distinct groups relating to ECM organization, collagen biosynthesis, and ECM degradation (**Figure S13**). In addition, clusters involving syndecan/non-integrin cell-surface signaling, integrin MAPK signaling, complement cascades, and toll-like receptor signaling were all enriched. Fibrinogens linked toll receptors and clot formation to ECM signaling, while FN1 again spanned categories of ECM organization and cell surface signaling by integrins and syndecans (**Figure S14**). LiCl extract analysis displayed the same ECM-related terms as in Azo but also captured TGFβ signaling (p-value=2.2E-02). Plotting relevant ECM-related terms, it is clear there is a significant degree of overlap between TGFβ signaling and ECM proteins involved in processes of extracellular structure and cellular adhesion (**Figure 4B**). In particular, THBS1 (thrombospondin-1), FN1 (fibronectin), and COL1A2 (collagen alpha-2(I) chain) appear to be focal points connecting these processes (**Figure 4B**).

### Bidirectional ECM–Myocyte Signaling in ICM

The bidirectional nature of ECM signaling with cells indicates that its composition likely plays a greater role in disease progression beyond deposition following injury and cardiomyocyte death. In ICM, the mechanical properties of ECM are altered,^47–49^ which may enhance pro-fibrotic TGFβ signaling and further drive fibrosis progression. This concept is supported by studies showing that engineered heart tissues derived from stem cells grown on ECM from hypertrophic hearts exhibited contractile dysfunction.^50^ Consistent with these findings, our data reveal alterations in both the ECM and the contractile machinery of cardiomyocytes. **Figure 5A** shows a schematic of the connective proteins linking ECM to internal contractile machinery via costameres,^51^ and **Figure 5B** shows the significant proteins in our experiment corresponding to these linkages. Costameres are similar to the focal adhesions cardiomyocytes have at their distal ends, but connect laterally to form sarcolemmal-cytoskeletal attachments.^47,51^ In addition, costameric proteins shown to interact with integrins such as kindlins (FERMT2/3) and parvins also significantly change at end-stage ICM. FERMT2 (also known as kindilin-2) is an essential component of vertebrate intercalated discs and is essential for cytoskeletal organization and membrane attachment during development. ^52^

**Figure 5.**
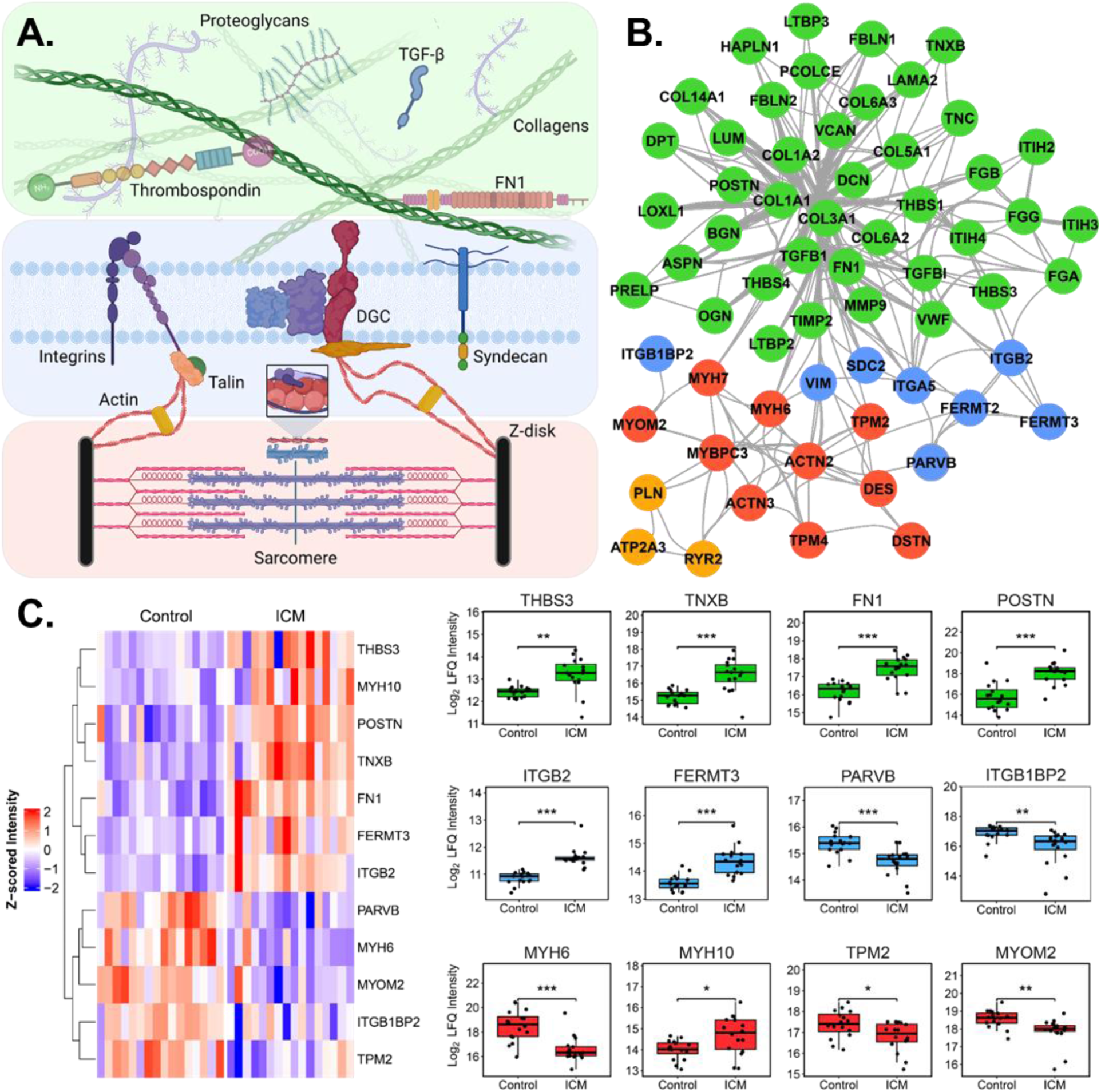
Bidirectional communication between ECM and cardiomyocytes contributes to ICM. **A)** A general scheme for interaction between ECM and cardiomyocyte contractile machinery is shown. At the sarcolemma, integrins and other costameric proteins transmit forces from ECM to cytoskeleton and then to sarcomere. **B)** Depiction of significantly altered protein groups from our data and the corresponding protein-protein interactions plotted in Cytoscape.^35^ **C)** Heat map and box plots of specific protein groups, demonstrating increased expression of ECM consistent with fibrosis, and changes to intracellular components of cardiomyocytes including fermitins, actinins, and myosin heavy-chain isoforms.

We observed a significant down regulation of kindlin-2 in ICM, consistent with previous studies in mice showing that deletion of this gene results in HF and premature death.^53^ Interestingly, loss of FERMT2 was contrasted by upregulation of FERMT3 and ITGB2, both of which are exclusively expressed in hematopoietic cells. The interaction of FERMT3/ITGB2 is particularly important in platelet aggregation and innate immune response,^54,55^ and is associated with unstable atherosclerotic plaques.^56^ We also observed the significant downregulation of muscle-specific integrin-binding protein melusin (ITGB1BP2), which protects the heart during ischemic injury via integrin-dependent activation of PI3K/AKT and ERK pathways.^57^ **Figure 5C** shows the specific proteins that change at the interconnected stages of ECM, costamere/sarcolemma, and sarcomere. These include the adhesive glycoproteins like fibronectin, thrombospondins and tenascins which all increase in ICM and components of the costamere like kindlins, parvins, and melusin. Intracellularly, myosin heavy chain 6 greatly decreases, as do tropomyosin-2 and myomesin-2, while myosin heavy chain 10 increases. Overall, there was a trend toward decreased expression of the intracellular components relative to the extracellular proteins. It is likely this is attributable at least in part to replacement fibrosis – cell death decreases the relative expression of intracellular components as ECM deposition takes the place of cells. Despite these contributions, interstitial fibrosis resulting in intracellular change is also very likely, given most patients in this study had no history of MI and had only experienced chronic ischemia.

## Discussion

Myocardial ischemia triggers the activation of cardiac fibroblasts stimulated by mechanical forces, growth factors, and cytokines to remodel the myocardium.^9,10,58^ Differentiation of these cells to myofibroblasts leads to pathological remodeling of myocardial ECM and subsequent cardiac fibrosis—key processes in the progression of ischemia to ICM and eventual HF.^8,58^ To identify differences in the composition of the ECM in end-stage failing ICM hearts versus nonfailing controls, we employed a sequential extraction to achieve maximal depth of coverage and focused primarily on analysis of the second extract, which relied on a photocleavable surfactant, Azo^24,59^, to solubilize core ECM proteins for analysis and quantitation by MS-based proteomics. For this study we compared differential protein abundance from 16 failing ICM tissues obtained during LVAD implantation to those from 16 region-matched nonfailing donors without a history of cardiovascular disease. Previous proteomics studies of human cardiac tissues have either focused on nonischemic disease^60^ or on the effect of LVAD implantation by using patient-matched pre- and post-operative samples.^61^

Given the inherent challenges in solubilizing ECM proteins owing to their insolubility, extensive crosslinking, and structural complexity, efforts have been dedicated to developing specialized extraction methods to extract the ECM proteins from heart tissues.^13,22,23^ For example, Barallobre-Barreiro and colleagues utilized sequential extraction with 0.5 mol/L sodium chloride (NaCl), 0.1% SDS, and 4 mol/L guanidine hydrochloride (guanidine-HCl) to decellularize and sequentially extract ECM proteins enabling the identification and analysis of 139 ECM proteins from the porcine myocardium subjected to ischemia/reperfusion injury.^13^ Similarly, de Castro Brás et al. developed the Texas 3-step decellularization protocol, which sequentially uses salt, detergent, and chaotropic agents to enrich for ECM components in cardiac tissue, facilitating proteomic analysis of the cardiac ECM.^22^ In this study, we used sequential extraction with the photocleavable surfactant, Azo, allowing fast tissue decellularization, efficient extraction and enrichment of ECM proteins, which dramatically expanded the coverage of ECM proteins and permitted the analysis of >315 ECM proteins from human failing ICM and nonfailing donor hearts. Our method has shown high reproducibility for label-free quantitation and enabled the identification of significantly altered proteins in the myocardium of end-stage ICM patients in comparison to the controls.

The most significantly differentially expressed protein in our data was the elastin and collagen crosslinking protein LOXL1 (lysyl oxidase homolog 1; adjusted p-value=3.57E-31; log2 fold change=2.6). LOXL1 was quantified in 12/16 ICM tissue samples using the Azo-based extraction and was absent in all nonfailing control samples, indicating a marked upregulation in failing hearts. Although its absolute intensity is relatively low, the detection and quantification of LOXL1 underscore the importance of our extraction method in capturing low-abundance proteins that may play vital roles in pathological remodeling. In animal models of hepatic and pulmonary fibrosis, LOXL1 has been shown to regulate TGFβ-induced fibrosis, and its knockdown inhibits fibrotic proliferation and reduces expression of type-I collagens and pro-fibrotic metalloproteinases.^40^ The substantial increase in LOXL1 observed in this study suggest a previously underappreciated role of this enzyme for myocardial stiffening and subsequent LV dysfunction in end-stage ICM, potentially through elastin crosslinking. Additionally, LOXL1 may contribute to collagen crosslinking, a process associated with increased myocardial interstitial fibrosis (MIF) and adverse clinical outcomes.^62,63^

We also observed a significant increase in the gelatinase MMP9 (matrix metalloproteinase-9, adjusted p-value=.026, log2 fold change=0.951), though to a lesser extent than LOXL1. MMP9 is known to degrade type-I collagen fibers, and the greater relative increase in crosslinking enzyme lends credence to the idea that MIF in HF involves crosslinking predominating over matrix degradation.^62^ Alongside these changes, several tissue inhibitors of metalloproteinases (TIMPs) and inter-alpha-trypsin inhibitors (ITIHs) were significantly up-regulated in ICM tissues. These proteins regulate ECM deposition by inhibiting degradation by metalloproteinases, and it is likely that the imbalanced expression of the proteases and their inhibitors contributes to the transition from ischemia to HF.^12,64^

Alongside crosslinking, collagen fiber composition and alignment vis-à-vis cardiomyocytes both affect the mechanical dysfunction of cardiac tissue in ICM.^65–68^ We observed an increased abundance of types I/III collagens and the expected decrease in I:III ratio associated with ICM, which is consistent with previous report.^66^ Elevation of these fibrillar collagens contributes significantly to cardiac fibrosis and can activate cardiac fibroblasts to create feedback loops of ECM deposition.^69,70^ Collagens I and III drive proliferation of undifferentiated cardiac fibroblasts whereas collagen VI (COL6A2, adjusted p-value=.0086, log2 fold change=1.02; COL6A3, adjusted p-value=0.0076, log2 fold change=0.932) potently induces myofibroblast differentiation.^71^ Mouse knockout of collagen VI promotes healing and reduces fibrosis, seemingly due to rapid myofibroblast apoptosis which prevents excessive matrix deposition.^72^ We also observed significant increases to the SLRPs asporin (ASPN), biglycan (BGN), decorin (DCN), lumican (LUM), and mimecan (OGN), which is known to bind collagens and regulate fibrillogenesis.^73^ SLRPs generally modulate collagen fibril formation and thus ECM integrity, tensile strength, and organization. In a mouse model of ICM, ASPN was one of the top differentially expressed genes and exhibited an anti-fibrotic effect by attenuating TGFβ signaling.^74^ DCN is also antifibrotic, slowing the rate of fibrillogenesis and blocking TGFβ-induced collagen deposition in three-dimensional collagen matrices.^75^ Contrary to this, LUM induces collagen I expression, as well as matrix remodeling molecules such as LOX, TGFβ, and MMP9.^15^ Further efforts into the PTMs of SLRPs including both N-linked glycosylation and GAG chain composition/sulfation is needed to elucidate their specific roles in ICM and HF.

TGFβ signaling is the central mediator of post-infarction cardiac repair; however, by stimulating ECM preservation and myofibroblast differentiation/activation, it also serves as a key contributor to fibrotic remodeling.^45^ Previous proteomics analysis of ECM in a porcine ischemia/reperfusion model identified a central role for TGF-β1 signaling in orchestrating early- and late-stage ECM remodeling.^13^ We observed significant increases to TGFβ1 (TGFB1) and TGFβ-related proteins including latent transforming growth factor binding proteins (LTBP1/3/4), which is known to play a critical role in the maintenance, extracellular secretion, and integrin-mediated activation of TGFβ.^76^ We also noted a decreased abundance of SMAD1, a molecule downstream of TGFβ activation, which has a documented antiapoptotic effect in cardiomyocytes.^77^ The fine detail of how ECM composition influences TGFβ activation remains unclear, but it is known that mechanical stiffening and fibrosis lower the activation threshold for TGFβ.^78^ The different isoforms of LTBPs show different matrix binding preferences, for example LTBP1 mainly interacts with fibronectin, mediated by heparan sulfate proteoglycans, and LTBP3/4 bind fibulins.^78^ Our experiment showed increased abundance of fibronectin and fibulins in failing ICM hearts, alongside other proteins that mediate the interaction between latent TGFβ and the ECM like small leucine-rich proteoglycans and integrins. These findings also support a TGFβ-driven feedback loop, in which ECM stiffening enhances mechanosensitive TGFβ, which in turn promotes further ECM deposition and mechanical remodeling of the matrix properties.^78,79^ Additional regulatory inputs, such as the renin-angiotensin-aldosterone system, further contribute to fibroblast activation and subsequent ECM deposition by stimulating TGFβ and related pathways.^80^ Beyond mechanical and systemic regulation, paracrine signaling by secretion of matricellular proteins is pivotal in mediating interactions between ECM and cells.^47^

In our experiment, we observed significant changes to adhesive glycoproteins and matricellular proteins in end-stage failing ICM hearts, including tenascins (TNC and TNXB), thrombospondins (THBS1/3/4), periostin (POSTN), and von Willebrand factor (VWF). Thrombospondin-1 is directly involved in TGFβ activation by forming an interaction with the TGF-β propeptide preventing formation of latent complexes, which increases the amount of the mature protein which can bind to receptors.^81^ Thrombospondins 1, 2 and 4 are known to play protective roles in the progression of cardiac diseases and have been explored as therapeutic targets.^82^ Recent mouse studies revealed that THBS3 destabilizes membranes by downregulating integrin.^83^ It was also shown that its expression was higher in the blood of patients with stable CAD compared with those who had experienced MI.^84^ For the first time, we have shown that THBS3 is upregulated significantly in human end-stage ICM, likely contributing to the disease progression in a similar manner. Overall, thrombospondins appear to be key modulators of integrin signaling and cardiac remodeling.

Although there is no standard definition for ICM, it can be considered LV dysfunction occurring with severe CAD and either history of MI or stenosis of >75% of left anterior descending artery/two or more coronary arteries.^5^ Improvements in patient survival following MI have resulted in a greater prevalence of ICM-induced HF.^85^ In spite of this, fewer than half (6/16) of the ICM patients in our study had a documented MI with the remainder diagnosed based on the severity of CAD. Even in the absence of MI, chronic ICM still results in profound myocardial interstitial fibrosis, though the underlying mechanism and pathophysiology of this outcome are unclear. In fact, Frangogiannis *et al*. have shown that demonstrated that the extent of interstitial fibrosis distinguishes irreversibly dysfunctional myocardium from tissue capable of functional recovery, suggesting chronic ICM progresses through ongoing fibrotic remodeling.^45,58,86^ Our data implicates TGFβ signaling as an important factor in end-stage HF from ICM, though the specific mechanisms of activation remain to be investigated. Previous studies suggested that acidity or hypoxia-induced signaling may contribute to TGFβ activation in ICM,^44^ and likely that the underlying cause of disease progression is dysregulation of TGF-β signaling itself. For example, we found SMAD1, a cytosolic protein that is phosphorylated in the canonical TGFβ pathway that protects against apoptosis in ischemia/reperfusion injury models,^77^ is downregulated in ICM tissues.^87^ There are ultimately many mechanisms that activate latent TGF-β, including the aforementioned THBS1,^81^ extracellular proteases,^88^ and mechanical strain.^78^ A feedback loop occurs in fibrosis when sufficiently stiff ECM reduces the threshold for TGF-β activation, which in turn results in myofibroblast activation and more ECM deposition.^78,89^ Our data suggest this process is possibly occurring in ICM, as we observe significant increases to both structural ECM components and many TGF-β related proteins in end-stage ICM tissues. Further studies are needed to elucidate how chronic ischemia drives myocardial matrix remodeling and interacts with dysregulated TGF-β signaling.

Overall, we have conducted a quantitative proteomics experiment to analyze the molecular changes, particularly in the ECM, of heart tissues from patients in end-stage HF from ICM. Our study revealed the drastic changes to ECM deposition in the ICM heart, including increased abundance of adhesive glycoproteins, matricellular proteins, and small leucine-rich proteoglycans. Notably, we found that THBS3 is significantly increased in ICM tissues and likely contributes to disease progression by destabilizing sarcolemmal integrin levels. This is consistent with our findings showing the dramatic increase in crosslinking protein LOXL1, suggesting altered mechanical properties of the matrix. Together, these findings point to a pathological interplay between TGFβ signaling and mechanosignaling via integrins as drivers of contractile dysfunction in the end-stage of ischemic disease. Altered protein expression at costameres, including integrins and their intracellular signaling partners such as kindlins and melusin, may further exacerbate the deleterious effects of increased fibrosis on cardiac function.

One limitation of our study is comparing ICM samples to only nonfailing hearts. Inclusion of other types of cardiomyopathies in the future would help identify ICM-specific ECM molecular signatures. Additionally, by using relatively small portions of LV tissue may not fully capture cardiac remodeling in ICM and thus may be susceptible to the effects of regional heterogeneity. Methodologically, although multi-step sequential extractions are widely used in ECM proteomics,^17,19,22,23,25^ they may complicate quantification due to overlap of proteins across fractions. Our group has recently reported a single-step extraction yields comparable results to two-step protocols, offering a potentially improved approach for future ECM quantification.^90^ Further investigation is needed to understand the origin of TGFβ dysregulation and its role fibrosis and their causal linkages to ischemia. Our data also revealed large-scale reduction in oxidative metabolism, and it would be prudent to investigate links between the metabolic phenotype of cardiomyocytes in ICM and the progression of interstitial fibrosis.

## Acknowledgements

This work was supported by NIH R01 HL109810 (to Y.G.). Y.G. also would like to acknowledge GM117058 and S10 OD018475. E.A.C. would like to acknowledge support from the NIH Chemistry-Biology Interface Training Program NIH T32GM008505. T.J.A. would like to acknowledge support from the Training Program in Molecular and Cellular Pharmacology, T32 GM008688-20. H.T.R. would like to acknowledge support from the National Heart, Lung, and Blood Institute of the NIH under Award Number T32HL007936 through the UW-Madison Cardiovascular Research Center.

