## Supporting Information for "Extracellular Matrix Alterations in Chronic Ischemic Cardiomyopathy Revealed by Quantitative Proteomics"

### Table of Contents

#### Supplementary Figures

Figure S1. Total ion chromatograms across extracts for all biological replicates...……S-3

Figure S2. Normalized intensities across extracts for all biological replicates..….…….S-4

Figure S3. Pearson correlation across biological groups and extracts …….…………..S-5

Figure S4. Enrichment of insoluble ECM ………..…...……………….…………….…S-6

Figure S5. Principal component analysis across biological groups and extracts ……....S-7

Figure S6. Differential expression and GO analysis of Extract 1...................................S-8

Figure S7. Box plots of significant ECM glycoproteins from Azo extract.……………S-9

Figure S8. Box plots of significant ECM regulators from Azo extract..…………...…S-10

Figure S9. Box plots of significant proteoglycans from Azo extract ……….………..S-11

Figure S10. Box plots of significant collagens from Azo extract ………..……...….…S-12

Figure S11. Clustered pathfindR enrichment plot using GO terms …………………...S-13

Figure S12. Upset plot of the representative GO terms from each cluster....................S-14

Figure S13. Clustered pathfindR enrichment plot using Reactome terms ………….....S-15

Figure S14. Upset plot of the representative Reactome terms from each cluster..........S-16

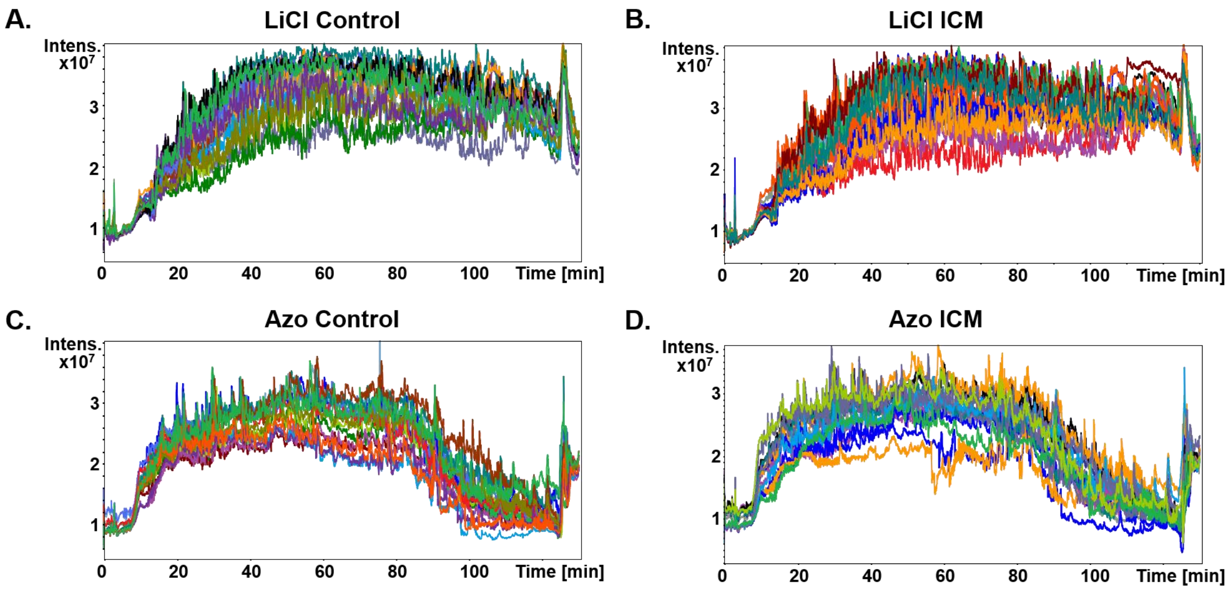


**Figure S1. Total ion chromatograms across extracts for all biological replicates.** Overlaid total ion chromatograms from bottom-up proteomic analysis for **(A-B)** LiCl and **(C-D)** Azo extracts in both cohorts (Control and ICM).

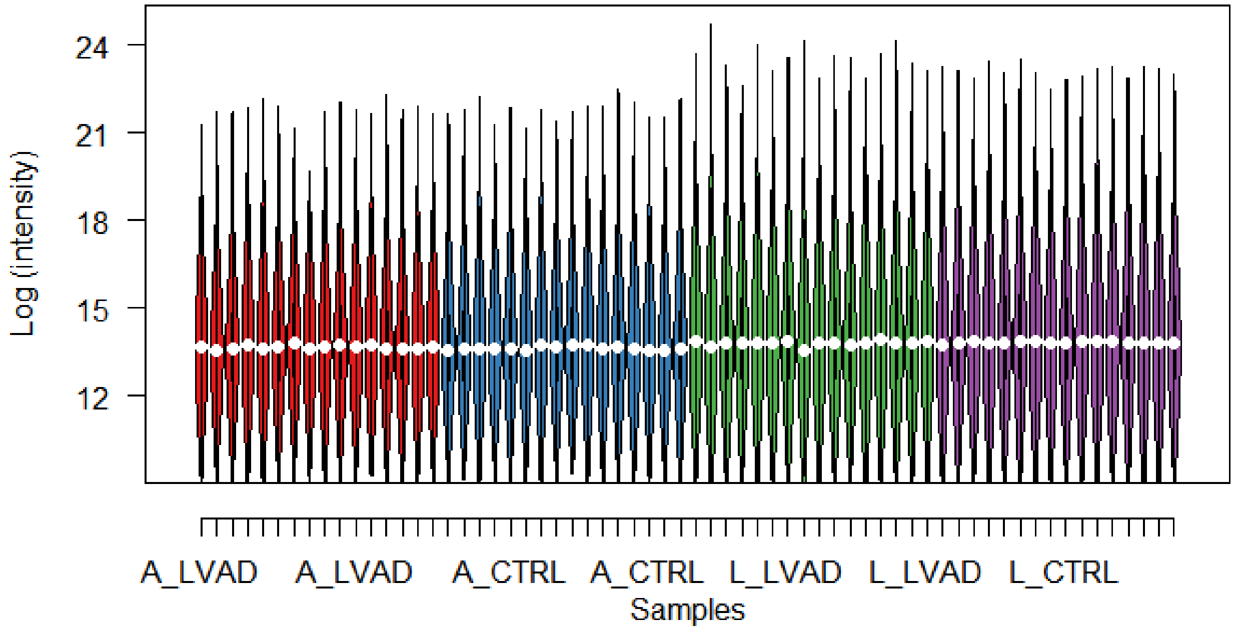


**Figure S2. Normalized intensities across extracts for all biological replicates.** To normalize data across conditions, sample intensity distributions were shifted to align the 15% quantile, and the shifted distributions are plotted here. Red and blue correspond to ICM and control samples from Extract 2, respectively, while green and purple refer to ICM and control samples from Extract 1.

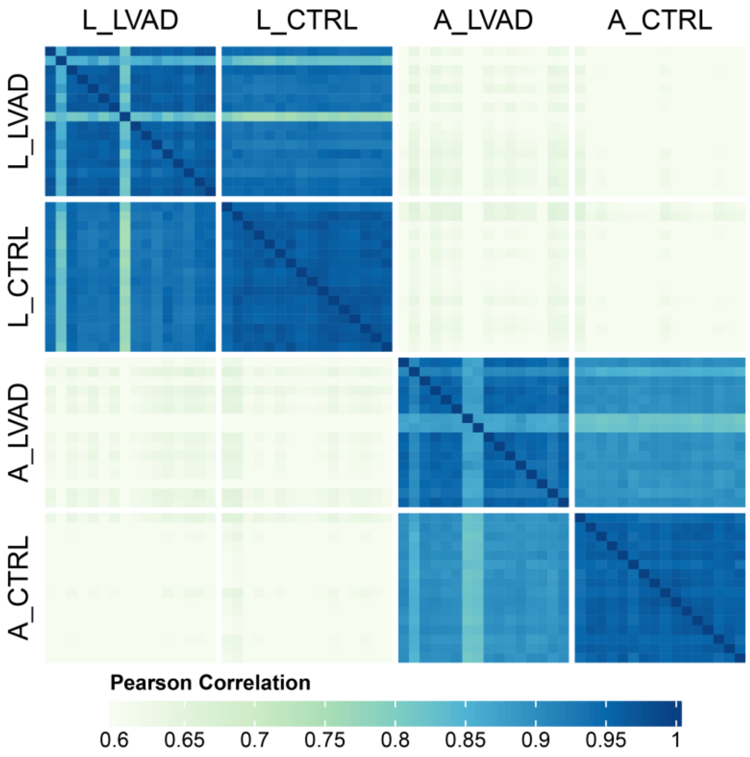


**Figure S3. Pearson correlation across biological groups and extracts.** Correlation matrix displaying color-coded Pearson correlation coefficients (r values) for all samples and conditions. The average Pearson correlation coefficient was r = 0.948 overall. For ICM samples, average r value for Extracts 1 and 2 respectively were 0.936 and 0.93, while controls were 0.966 and 0.96.

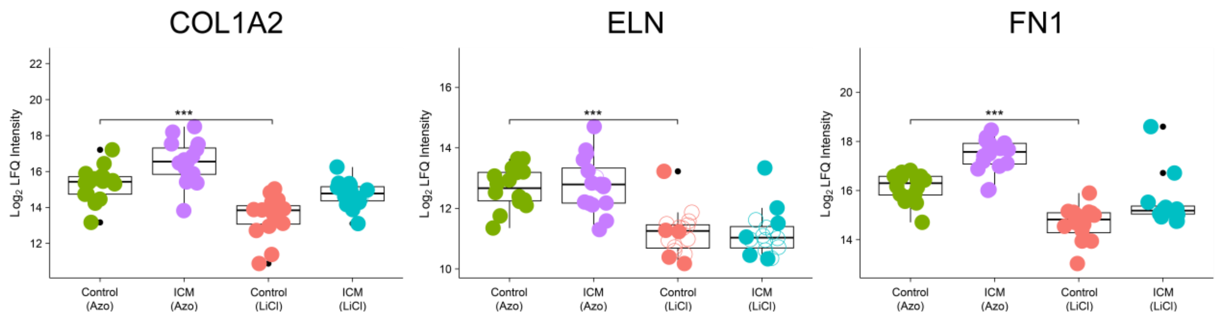


**Figure S4. Enrichment of insoluble ECM.** Box plots showing the $\log_2$ transformed intensities of COL1A2, ELN, and FN1 identified in both extracts, demonstrating use of Azo significantly enriched for ECM proteins.

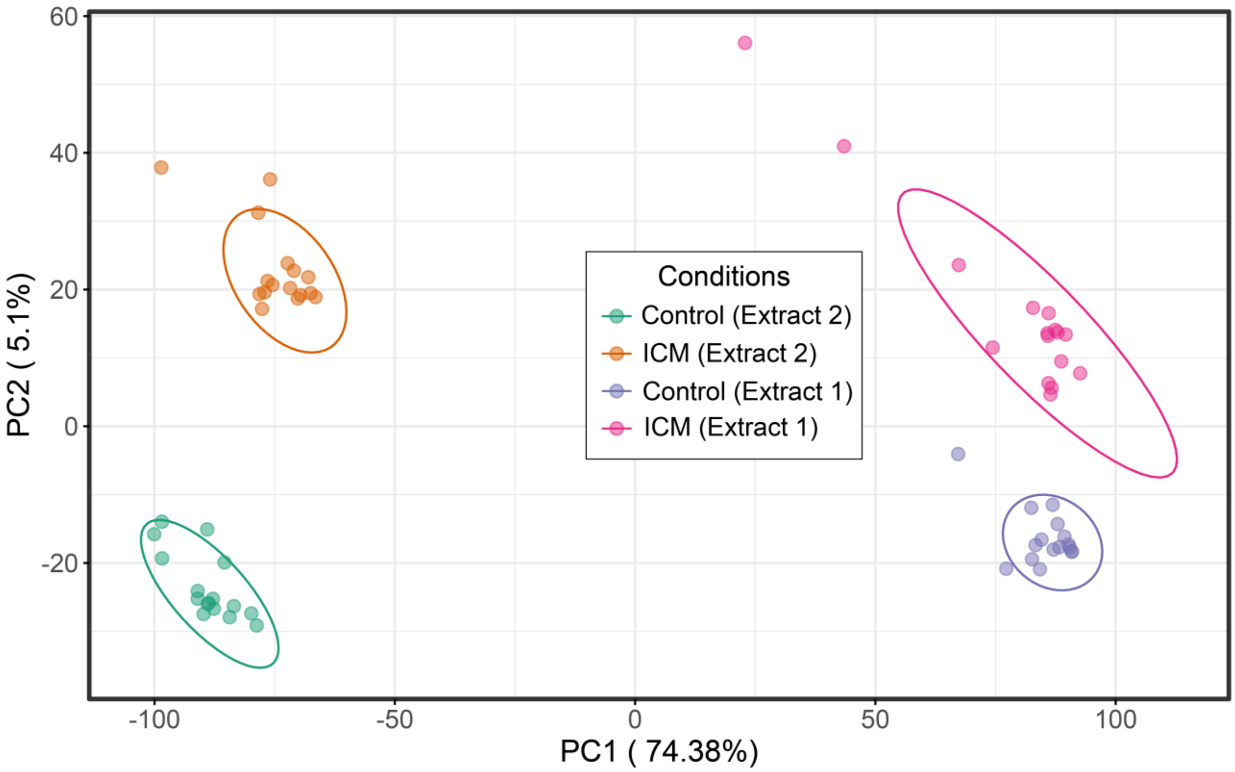


**Figure S5. Principal component analysis across biological groups and extracts.** Plot of principal component analysis for all conditions.

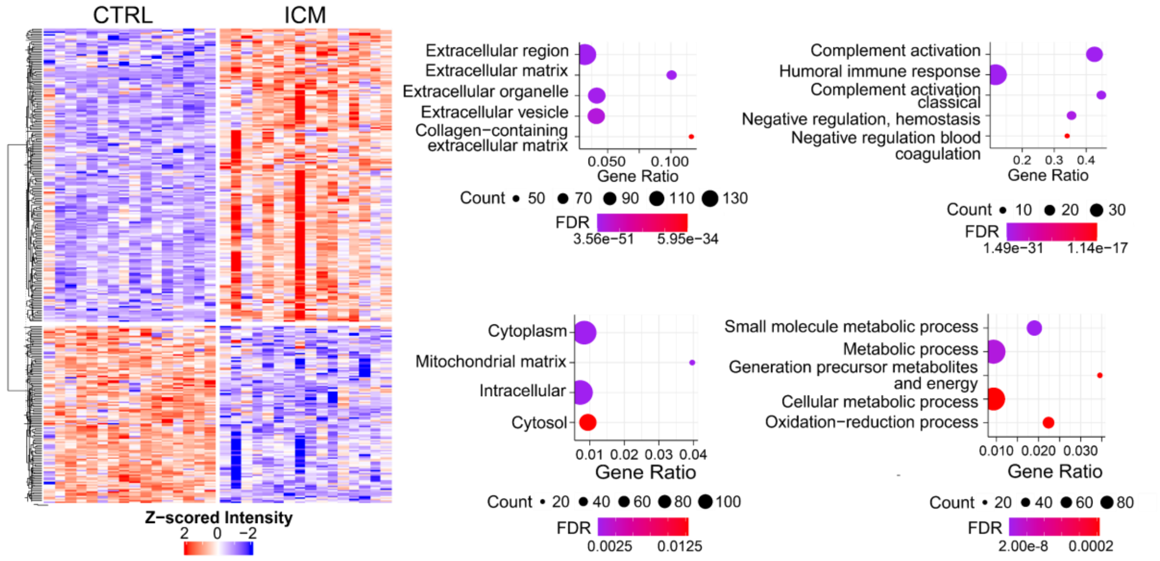


**Figure S6. Differential expression and GO analysis of Extract 1.** Heatmap of most significantly changed (by adjusted p-value) protein groups in Extract 1 and corresponding GO analyses of clusters. The first column of GO plots are cellular compartments and the second are biological processes.

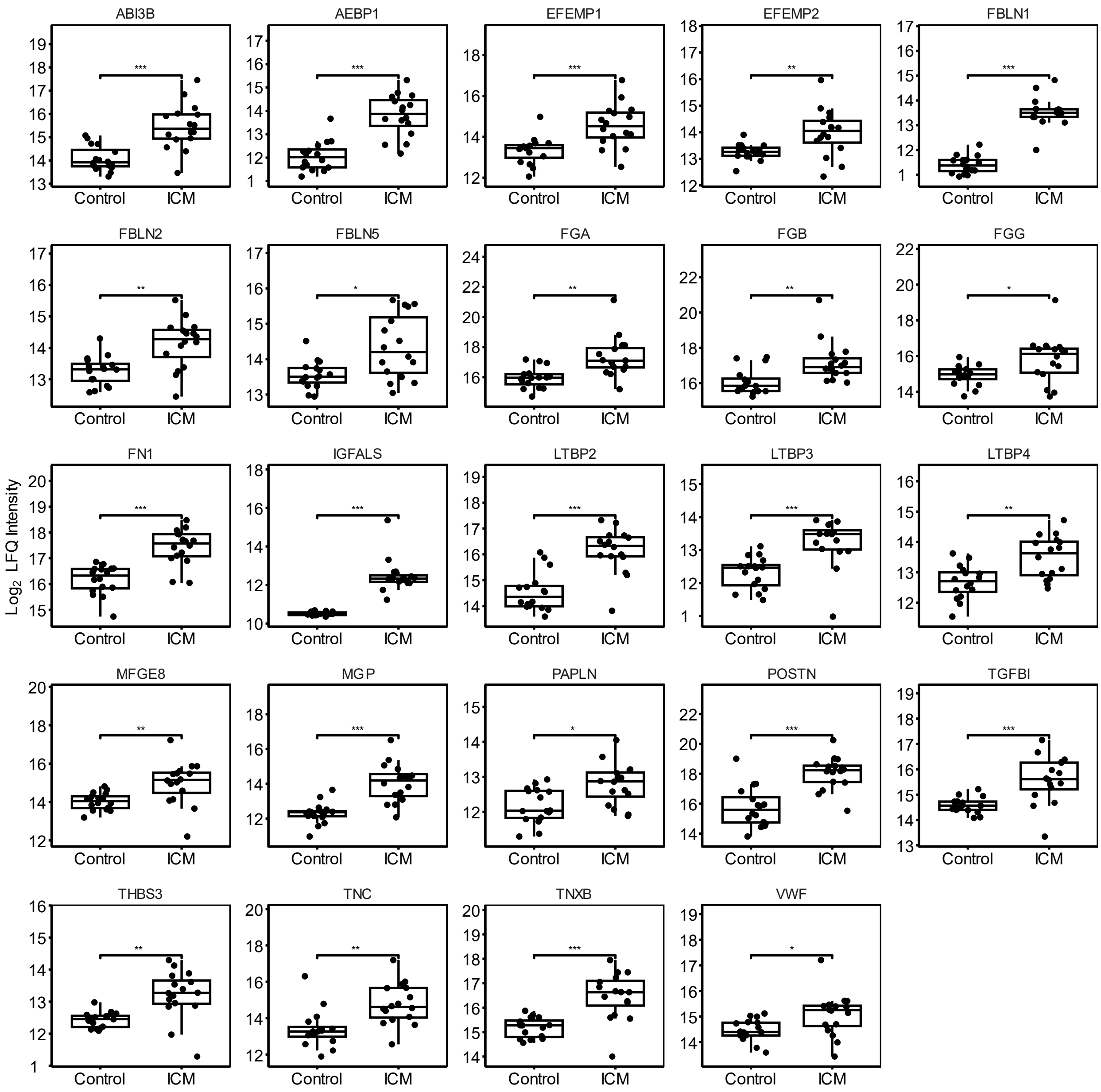


**Figure S7. Box plots of significant ECM glycoproteins from Azo extract.** Box plots displaying significant glycoproteins, with all replicates shown. P-values adjusted by independent hypothesis weighting are plotted as follows: * ≤ 0.05, ** ⩽ 0.01, *** ⩽ 0.001.

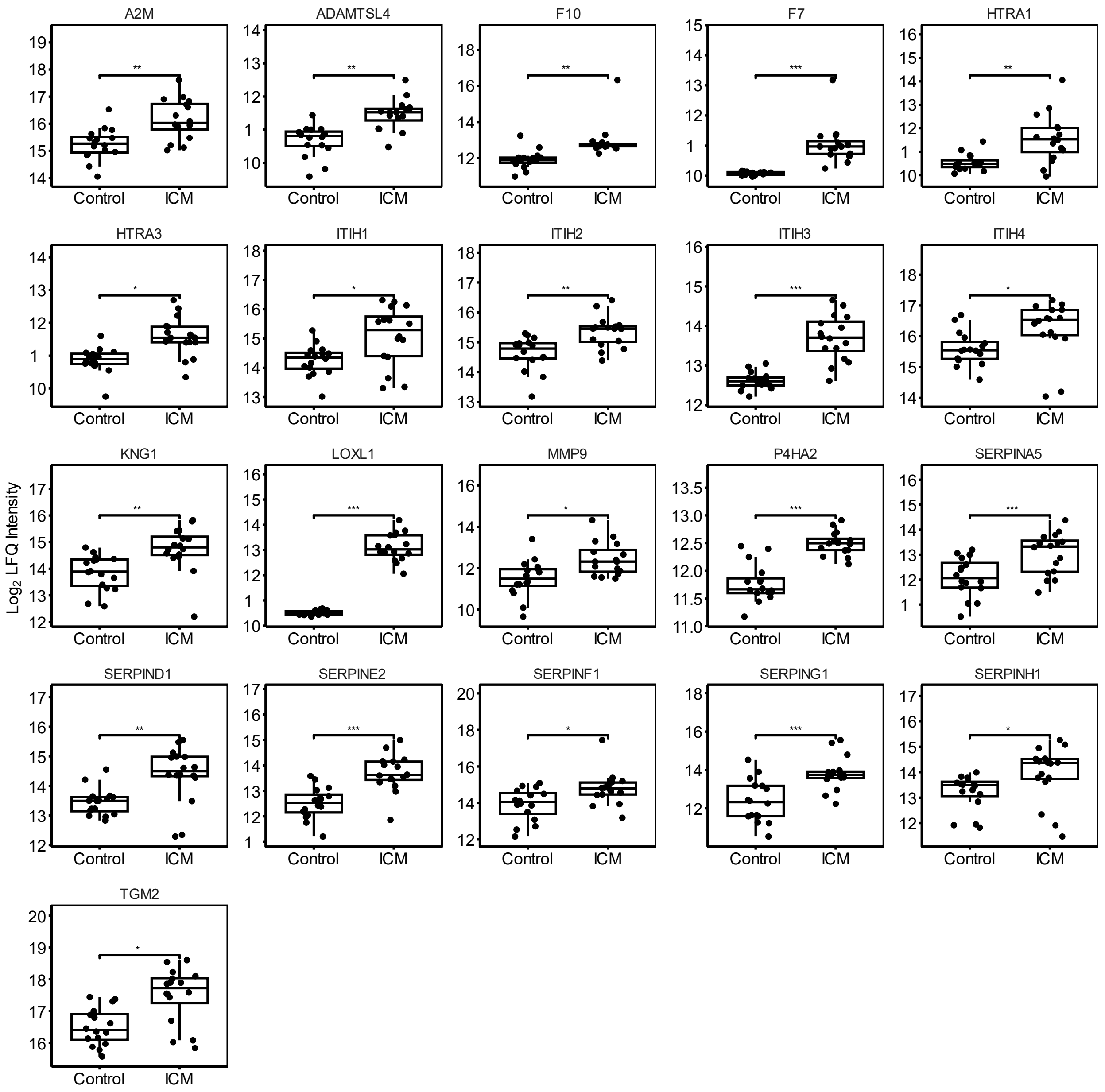


**Figure S8. Box plots of significant ECM regulators from Azo extract.** Box plots displaying significant regulators, with all replicates shown. P-values adjusted by independent hypothesis weighting are plotted as follows: * ≤ 0.05, ** ≤ 0.01, *** ≤ 0.001.

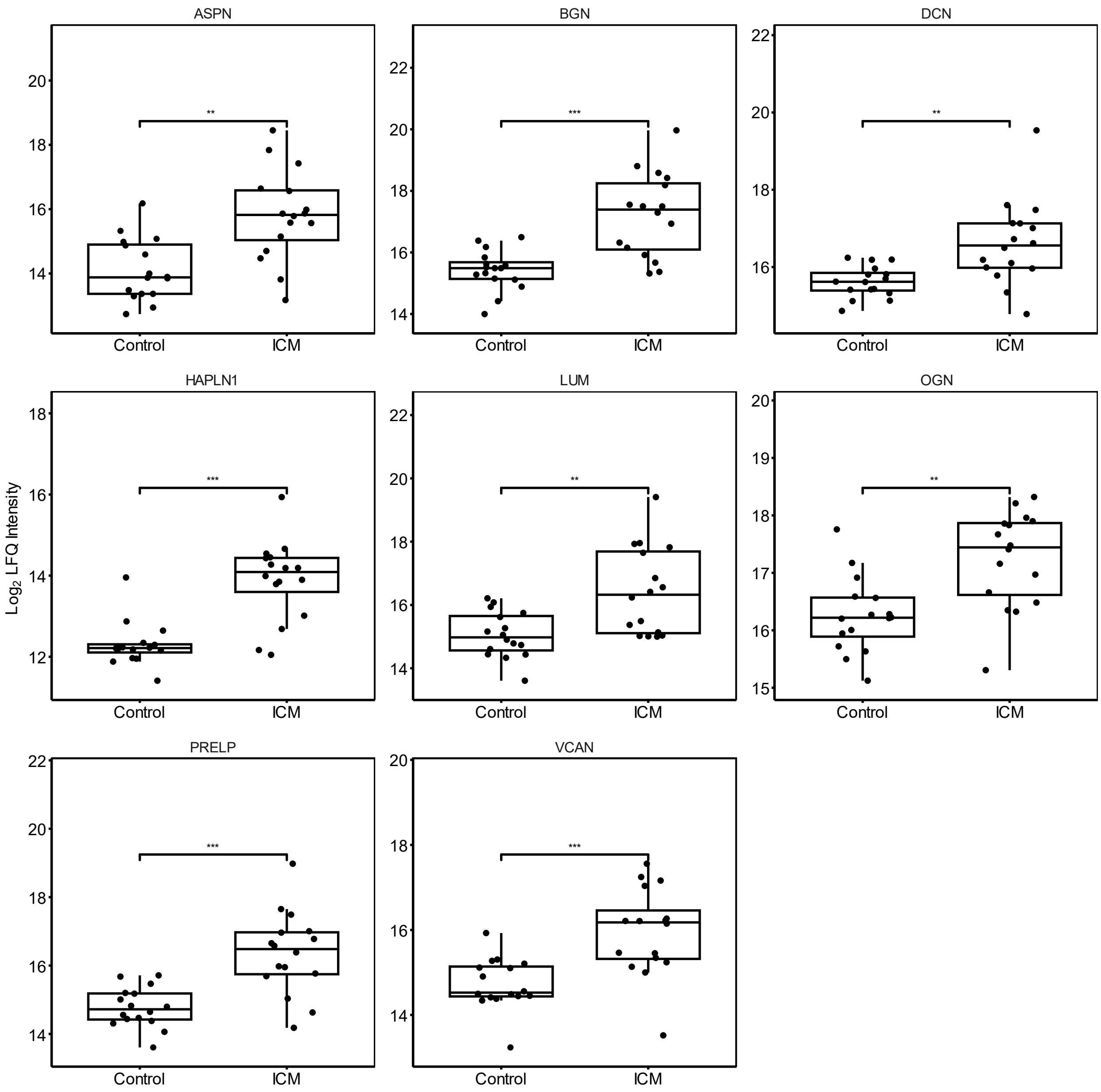


**Figure S9. Box plots of significant proteoglycans from Azo extract.** Box plots displaying significant proteoglycans, with all replicates shown. P-values adjusted by independent hypothesis weighting are plotted as follows: * ≤ 0.05, ** ≤ 0.01, *** ≤ 0.001.

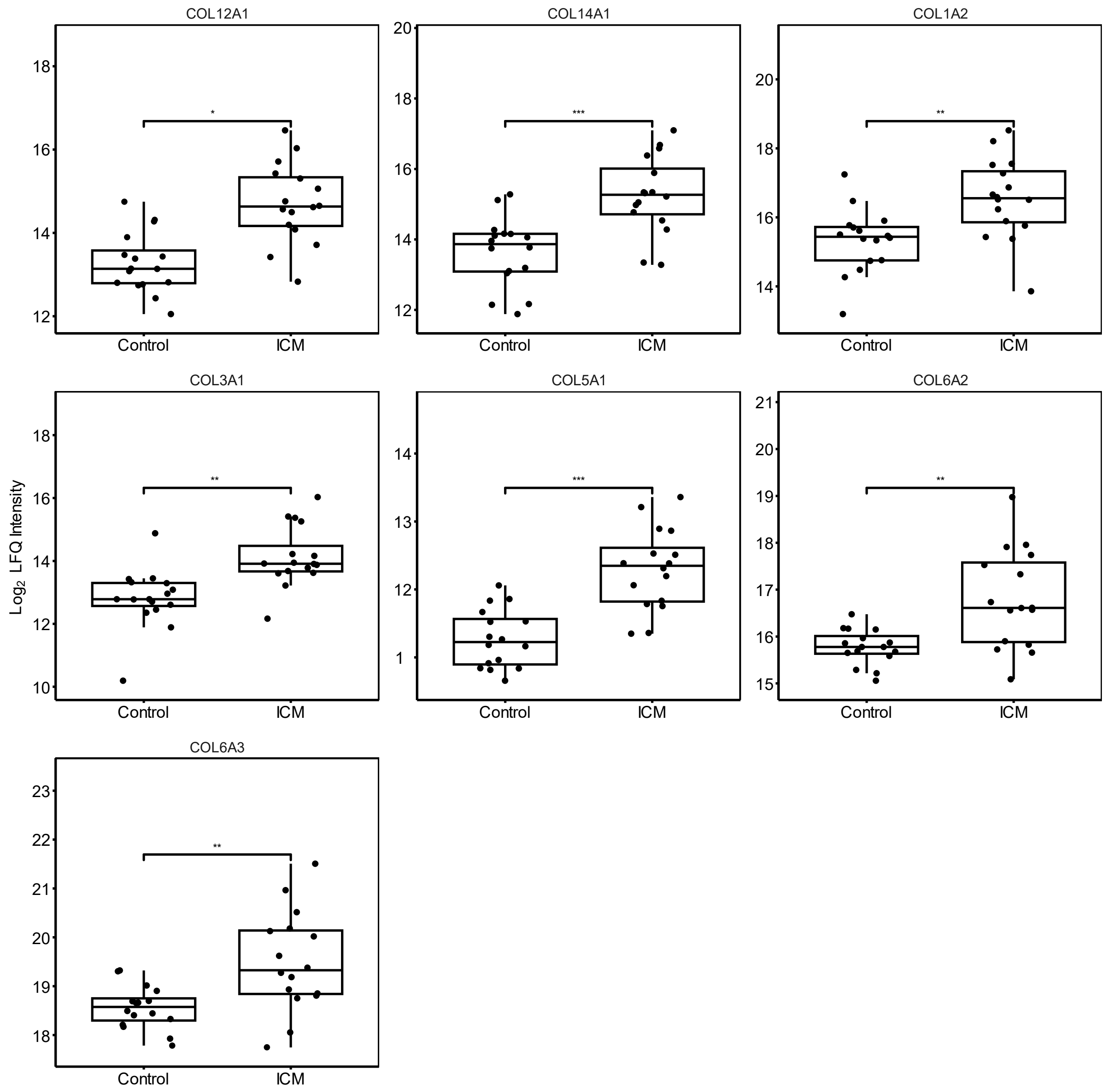


**Figure S10. Box plots of significant collagens from Azo extract.** Box plots displaying significant collagens, with all replicates shown. P-values adjusted by independent hypothesis weighting are plotted as follows: * ≤ 0.05, ** ≤ 0.01, *** ≤ 0.001.

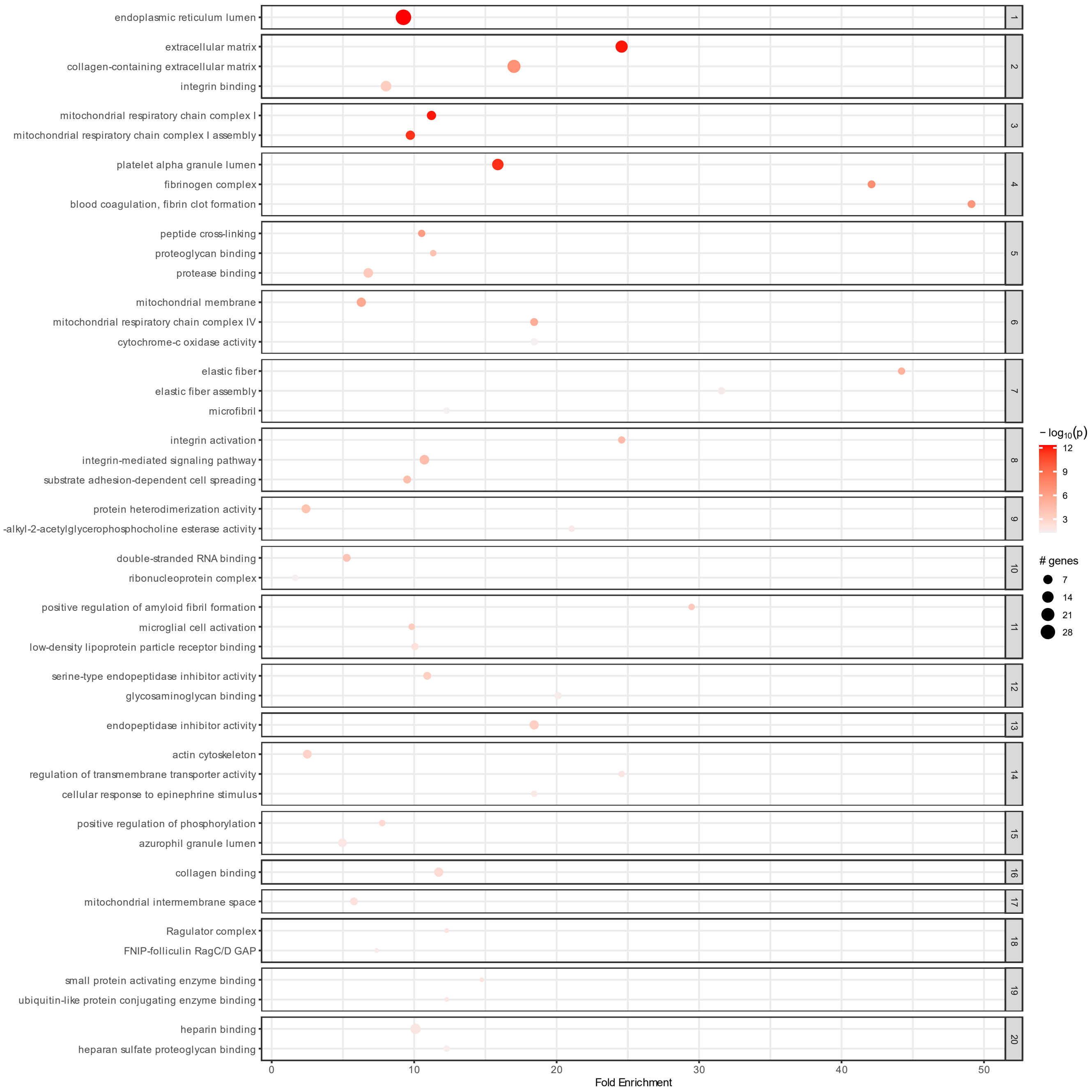


**Figure S11. Clustered pathfindR enrichment plot using GO terms.** Enriched terms from all GO categories (CC, MF, and BP) were hierarchically clustered. The top 20 clusters according to lowest enrichment p-values were plotted with a maximum of the three top terms per cluster. The first term in each cluster is designated as the representative term of its group. Dot size corresponds to the total number of proteins associated with the term, and color corresponds to lowest enrichment p-value for the term.

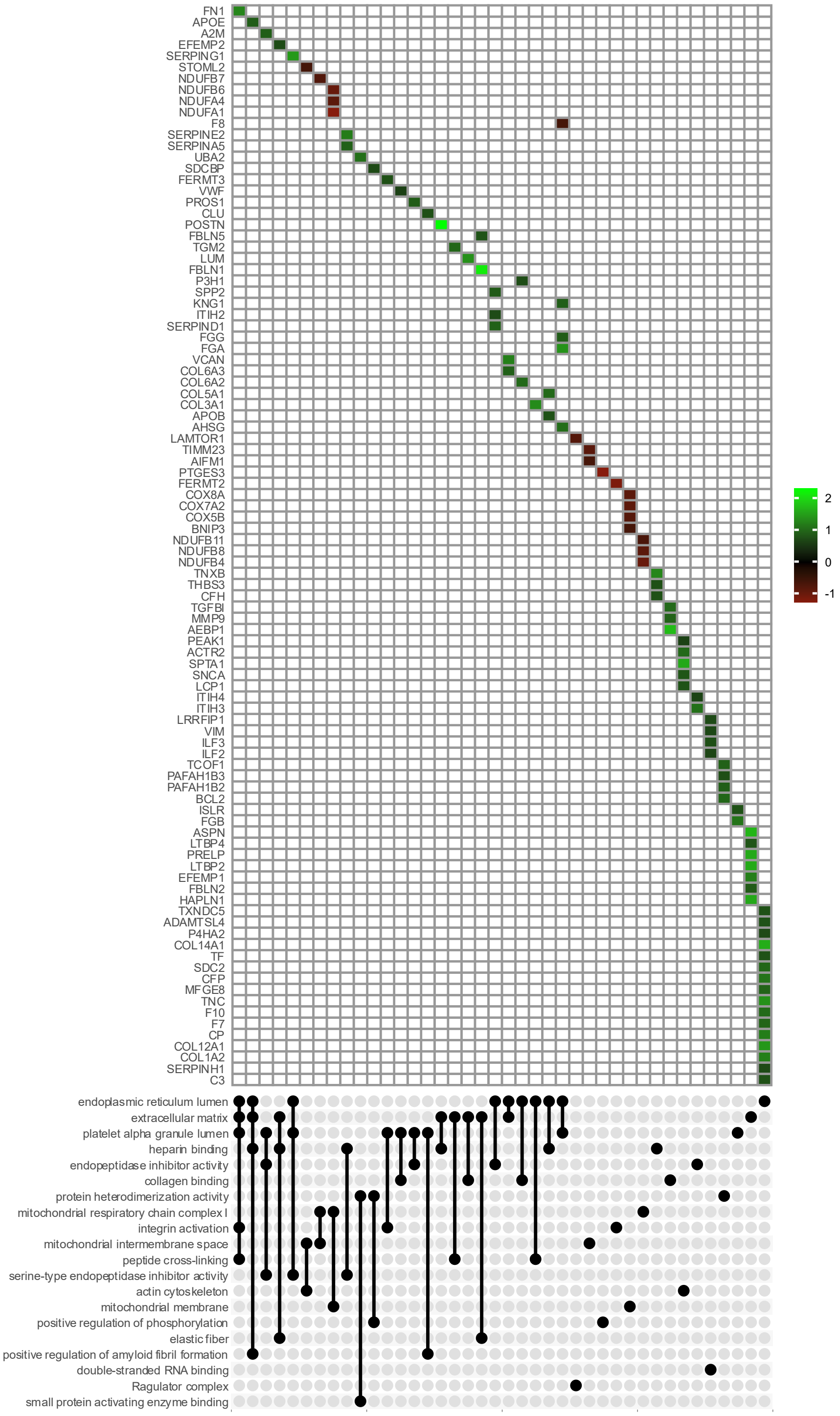


**Figure S12. Upset plot of the representative GO terms from each cluster.** Intersections of representative enriched GO terms from clustered pathfinder output were plotted for each protein in the intersection. Color corresponds to the $\log_2$ fold change ratio for the protein (control vs. ICM).

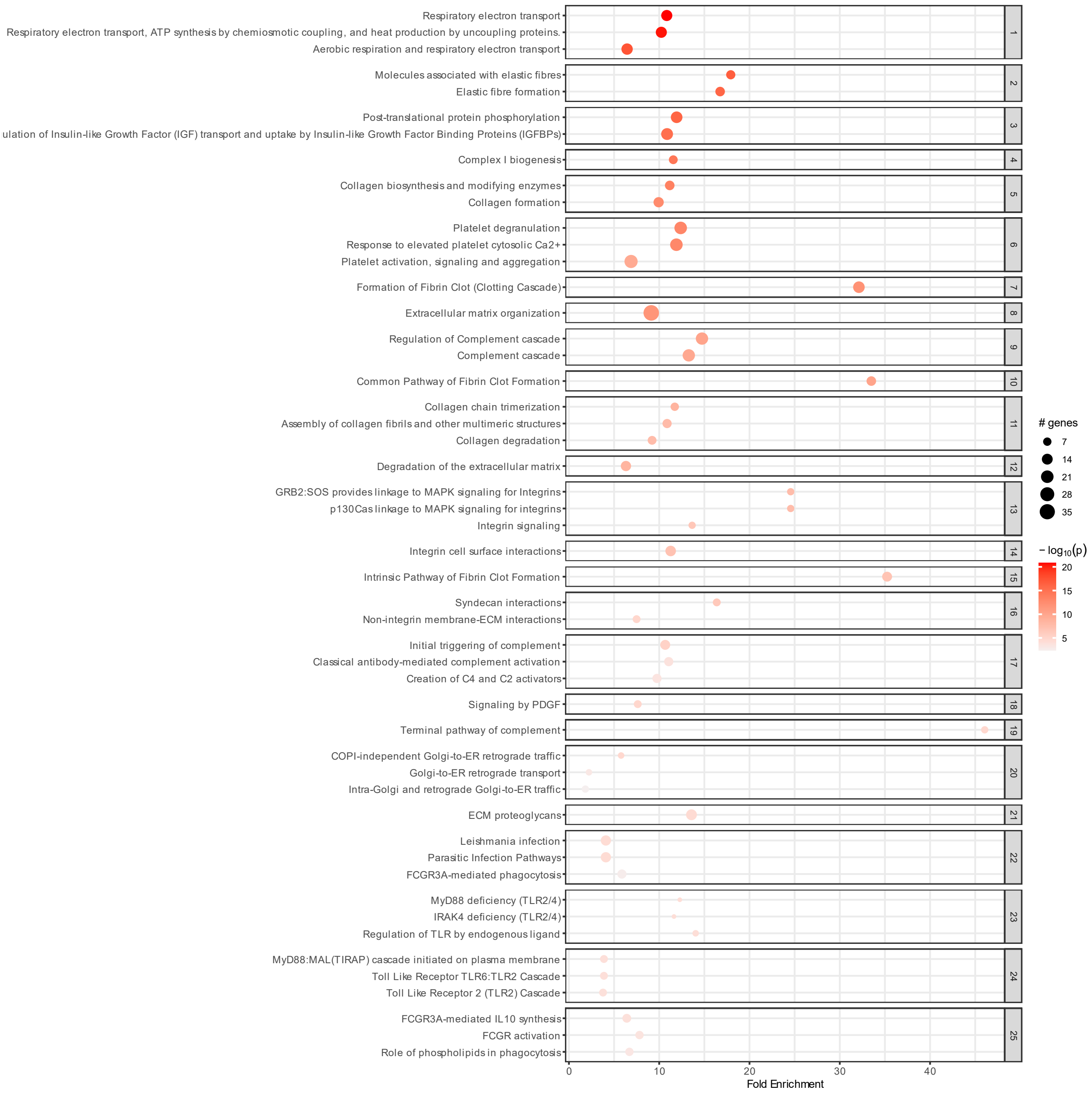


**Figure S13. Clustered pathfindR enrichment plot using Reactome terms.** Enriched terms from Reactome were hierarchically clustered. The top 25 clusters according to lowest enrichment p-values were plotted with a maximum of the three top terms per cluster. The first term in each cluster is designated as the representative term of its group Dot size corresponds to the total number of proteins associated with the term, and color corresponds to lowest enrichment p-value for the term.

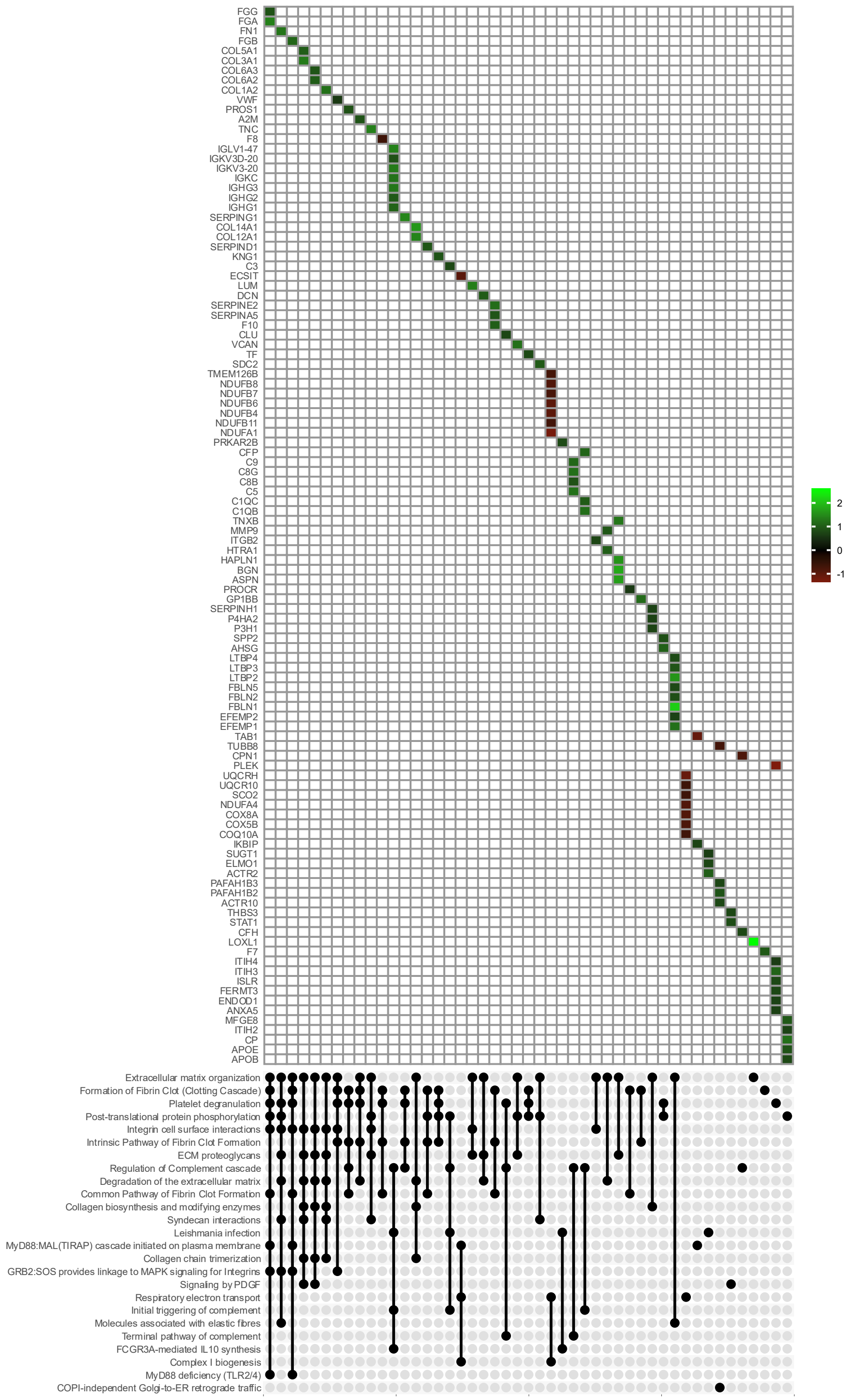


**Figure S14. Upset plot of the representative Reactome terms from each cluster.** Intersections of representative enriched Reactome terms from clustered pathfinder output were plotted for each protein in the intersection. Color corresponds to the $log_2$ fold change ratio for the protein (control vs. ICM).
